# LC3 subfamily in cardiolipin-mediated mitophagy: A comparison of the LC3A, LC3B and LC3C homologs

**DOI:** 10.1101/2020.07.14.202812

**Authors:** Marina N. Iriondo, Asier Etxaniz, Yaiza R. Varela, Javier H. Hervás, L. Ruth Montes, Félix M. Goñi, Alicia Alonso

**Author notes:** Corresponding autor. These two authors contributed to a similar extent to the studies described in this paper (Marina N. Iriondo, Asier Etxaniz).

## Abstract

Among the described indicators of mitochondrial damage, externalization of the phospholipid cardiolipin (CL) to the outer mitochondrial membrane has been proposed to trigger mitophagy, acting as a signal for binding the autophagy protein LC3B. However, the behavior of the other LC3 subfamily members has not been explored yet. In the present contribution, a comparative analysis of the interaction of LC3B, LC3A and LC3C with CL-containing model membranes, as well as their ability to translocate to mitochondria was assessed. Binding of LC3A to CL was higher than that of LC3B, and both proteins showed a similar ability to colocalize with mitochondria upon induction of CL externalization by rotenone in HeLa-NDPK-D or SH-SY5Y cells. Two residues located in the N-terminal region of LC3A were shown to be important for its recognition of damaged mitochondria. Moreover, the *in vitro* results suggested a possible role of LC3A, but not of LC3B, in oxidized-CL recognition as a counterweight to excessive apoptosis activation. In the case of LC3C, even if this protein showed a higher binding than LC3B or LC3A to CL, binding was less specific, and colocalization of LC3C with mitochondria was not rotenone-dependent. These results suggest that, at variance with LC3A, LC3C does not participate in the rotenone-CL mitophagy mechanism. The data support the notion that the various LC3/GABARAP family members might play different roles during autophagy initiation, identifying LC3A as a novel stakeholder in CL-mediated mitophagy.

## Introduction

Mitochondria are essential organelles for energy transduction in the eukaryotic cell. Mitochondrial dysfunctions have been associated with a wide number of pathological conditions, including neurodegenerative diseases, myopathies, and cancer [**1**]. Cells have developed different quality control systems in order to maintain an optimal mitochondrial network, including processes that allow the removal of damaged or superfluous mitochondria without causing cell death [**2**]. Among them, the selective degradation of mitochondria via macroautophagy is termed mitophagy.

Macroautophagy is characterized by the formation of a double-membrane structure around the cellular components to be degraded, the autophagosome (AP). The AP then fuses with lysosomes resulting in the degradation of contents by the lysosomal hydrolases. In this highly conserved mechanism, a plethora of signals and proteins are involved, making it exceedingly complex [**3**]. Among the different proteins involved in autophagy, the LC3/GABARAP protein family has attracted the attention of researchers [**4**]. This family is composed of at least six orthologs of the yeast protein Atg8, divided into two subfamilies, LC3 and GABARAP [**5**]. As in the case of Atg8, all LC3/GABARAP family members are found in both layers of the crescent autophagosomes [**6**], bound to phosphatidylethanolamine (PE) via a conserved C-terminal glycine due to the coordinated action of two ubiquitin-like systems [**7**]. This protein family is involved not only in membrane elongation, closure and fusion with lysosomes [**8,9**] but also in cargo recruitment [**10**].

Recognition of damaged or superfluous mitochondria is a key event in mitophagy. Many different indicators of mitochondrial damage are known. The PINK1/parkin axis, recognized as the main regulator of mitophagy [**11**], is characterised by the tagging of damaged mitochondria with ubiquitin chains that trigger their selective autophagy. This process requires the participation of different proteins such as optineurin and NDP52, which act as adaptors. They are able not only to recognize the ubiquitin chain on the mitochondrial surface but also to bind LC3B in the AP membrane [**12**]. However, a growing body of evidence has shown that this pathway is not the only relevant one. Many mitophagy receptors, such as FUNDC1, NIX, BINP3, Bcl2-L-13, AMBRA1 or FKBP8, that are found in the outer mitochondrial membrane (OMM), have shown the ability to recruit directly LC3/GABARAP proteins to mediate mitochondrial degradation, without ubiquitin signalling [**13**]. The majority of these receptors have a LC3 interacting motif (LIR) that allows their interaction with LC3/GABARAP [**14**]. Lipids, such as ceramide [**15**] or cardiolipin [**16**], can also serve as damaged mitochondrial signals/receptors.

The mitophagy mechanism mediated by the phospholipid cardiolipin (CL) is particularly intriguing. CL is essential for mitochondrial function and it is localized in the inner mitochondrial membrane (IMM) in healthy mitochondria [**17**]. In 2013, Chu et al. [**16**] found out that rotenone and other pro-mitophagic stimuli cause CL externalization to OMM, in turn inducing mitophagy. Furthermore, inhibition of CL synthesis or of CL externalization decreased the delivery of mitochondria to AP. Therefore, those authors proposed that CL would play a role in mitophagy; its externalization to the OMM upon mitochondrial injury would act as a signal for LC3 to remove damaged mitochondria in neuronal cells, preventing CL oxidation and accumulation of proapoptotic signals [**16**]. Recent studies have demonstrated that CL-externalization takes place also *in vivo*. Traumatic Brain Injury (TBI) model animals externalize CL to the OMM to a similar extent than rotenone-exposed mitochondria, thus underlining the importance of CL for mitophagy induction during the early response to this injury in human and rat brain [**18**]. Moreover, it is known that taffazin-deficiency related perturbations in CL remodeling related to Barth syndrome cause defective mitophagosome biogenesis [**19**]. In addition, it seems that externalized CL could interact with alpha-synuclein [**20**], a protein associated with Parkinson’s disease. For these reasons, studying this mechanism of CL-mediated mitophagy could lead to interpreting some neurodegenerative diseases as mitophagy-related diseases [**21**].

Previous studies from our laboratory [**22**] explored the molecular basis of LC3B-CL interaction, showing that LC3B was able to interact preferentially with CL over other dianionic lipids, and that after protein binding to CL-enriched membranes the C-terminus of LC3B remained exposed to the hydrophilic environment. This showed, as Chu et al. [**16**] had proposed, a mode of interaction between LC3B and CL that would allow the N-terminal region of LC3B to recognize damaged mitochondria, while its C-terminal region would target the autophagosomal membrane. Thus, CL appeared to serve as a specific mitophagy receptor for LC3B, ultimately leading to the removal of damaged mitochondria. However, no comparative study was available on the CL- and mitochondria-binding abilities of the LC3 subfamily members LC3A, LC3B, and LC3C. The data in this paper show the different behaviour of the LC3 subfamily members in the selection of damaged mitochondria as autophagosomal cargo, and describe LC3A as a LC3/GABARAP family member capable of recognizing externalized CL, and mediating mitophagy.

## Results

### The highly similar, predicted CL interaction regions of LC3 subfamily members suggest similar functions

In mammals a variety of well conserved Atg8 orthologs can be found (**Fig. 1**), suggesting some degree of specialization of the various family members. LC3/GABARAP proteins share a similar structure consisting of two amino-terminal α-helices and a ubiquitin-like (UBL) core (**Fig. 1A**). However, despite the extensive structural similarities, it has been shown that, while the first α-helix of LC3 subfamily is strongly basic, in the GABARAP subfamily this region is acidic [**23**] (**Fig. 1 A, B**). These differences are relevant because it has been proposed that the N-terminal region of these proteins might be important for specific functions during autophagy, playing an important role in protein-protein interactions, lipid-protein interactions, or via post-translational modifications [**24**].

**Figure 1.**
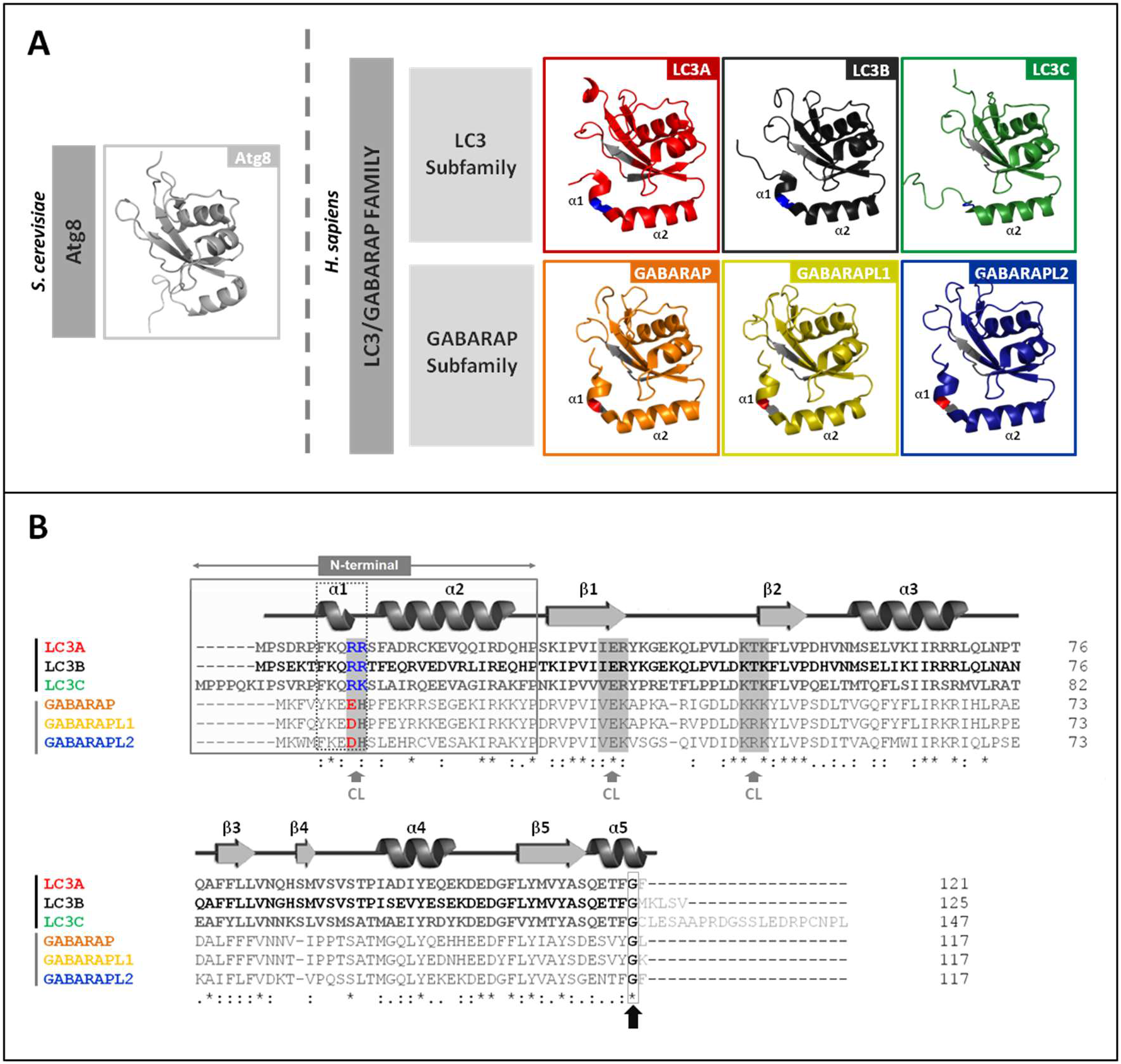
LC3/GABARAP family members and their interaction with CL. (**A**) Schematic representation of the 3D structures of yeast Atg8 and of each LC3/GABARAP family homologue in solution, displayed with PyMOL. PDB: Atg8 (2KQ7), PDB: LC3A (5CX3), LC3B (2ZJD), LC3C (2NCN), GABARAP (1GNU), GABARAPL1 (5LXI) and GABARAPL2 (4CO7). The thermodynamically preferred residues for cardiolipin binding on LC3B, identified by docking analysis [**16**], and their equivalents in the other LC3/GABARAP family members are highlighted in grey (See Fig 1B). Among them, the two residues in the α1 N-terminal region proposed to be essential for the interaction, and their equivalents in the LC3 subfamily, are colored in blue (positively charged). The amino acid corresponding to R10 of LC3B is a negatively charged residue in the GABARAP subfamily, colored in red (See Fig 1B). (**B**) Sequence alignment of the human orthologs of Atg8 obtained with Clustal W. LC3A (Q9GZQ8), LC3B (Q9GZQ8), LC3C (Q9BXW4), GABARAP (O95166), GABARAPL1 (Q9H0R8) and GABARAPL2 (P60520). The secondary structure elements of LC3B (PDBsum, 2ZJD) are indicated above the alignment as an example. The N-terminal and α-helix 1 (α1) of the two subfamilies are boxed. As in Figure 1A, the thermodynamically preferred residues for cardiolipin binding of LC3B, and their equivalents in the other members of the LC3/GABARAP family are highlighted in grey (grey arrow). The two positively charged α1 N-terminal residues proposed to be essential for the interaction of LC3B with CL and their equivalents are colored. In blue: positively charged. In red: negatively charged. The C-terminal glycine (black arrow) is conserved in all the LC3/GABARAP family members.

Previous studies from this laboratory have focused on the potential different roles of these two subfamilies, both in cargo recognition [**22**] and during autophagosome elongation [**25**]. Regarding cargo recognition, several aspects of the mitophagic mechanism initiated by CL externalization remain obscure. Previous work from this laboratory [**22**] compared the behaviour of LC3B with that of GABARAP subfamily members. The study explored the differences in residues putatively involved in LC3B interaction with CL (**Fig. 1B**). *In vitro*, the GABARAP subfamily members showed less binding to CL than LC3B and this corresponded with the absence of translocation to mitochondria in cells.

Taking into account the similarity between the predicted CL-interacting residues in LC3B and those in LC3A and LC3C (**Fig. 1B**), it could be assumed that all three LC3 subfamily members would be involved in CL recognition. This hypothesis was experimentally tested as follows.

### LC3A and LC3C, as well as LC3B, interact with CL-containing model membranes

To determine whether members of the LC3 subfamily other than LC3B were also able to bind CL-enriched model membranes, the binding of LC3A and LC3C to large unilamellar vesicles (LUV) containing CL was measured. For this purpose, a vesicle flotation assay was performed [**22**], in which protein association with membranes was assessed by the protein ability to float together with the vesicles after equilibrium in sucrose-gradient centrifugation (**Fig. 2A**). CL concentration in these experiments intended to mimic the one of the interaction zones formed in the OMM after CL exposure, and it was based on the CL levels used in similar studies with other proteins [**20,26,27**].

**Figure 2.**
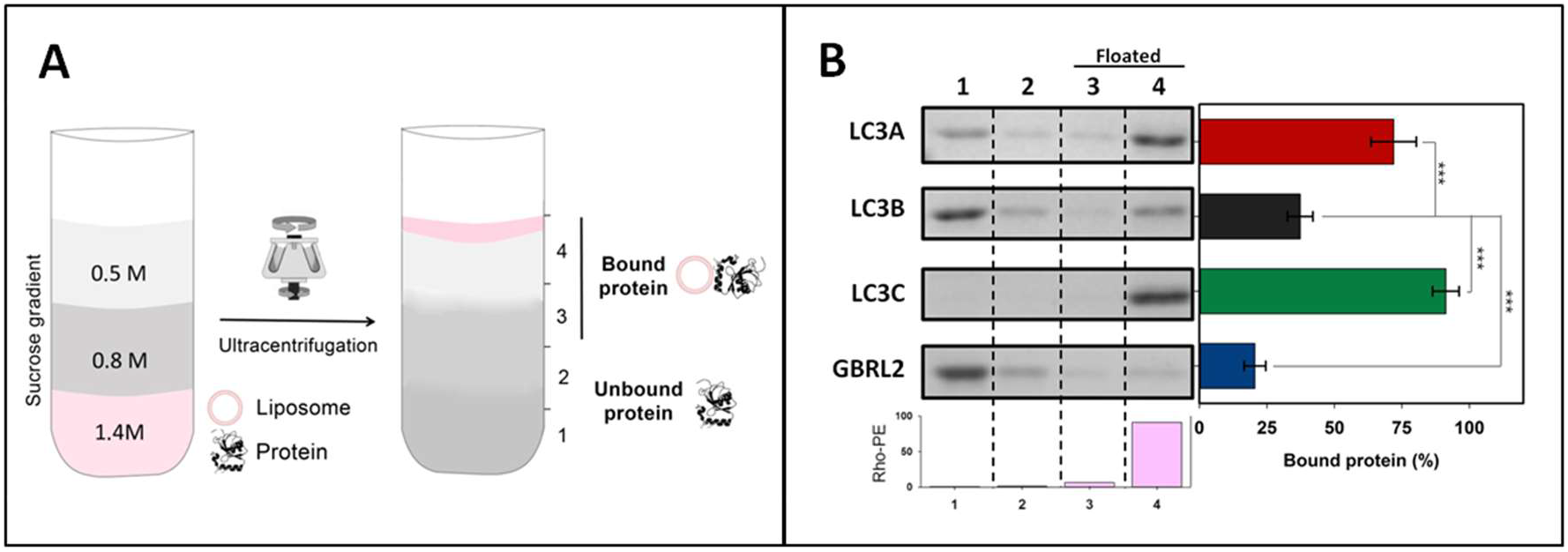
LC3A and LC3C, as well as LC3B, interact with CL-containing model membranes. Interaction of LC3/GABARAP proteins with CL-containing membranes measured by a vesicle flotation assay. (**A**) Representative scheme of a flotation assay in a sucrose-density gradient. Proteins (10 μM) were incubated with rhodamine-PE (Rho-PE)-labeled LUVs (3 mM) and subsequently mixed within the 1.4 M (final concentration) layer of a discontinuous sucrose gradient. After ultracentrifugation, four equal-volume fractions were collected, 1-4, starting from the bottom; protein found in fractions 3+4 was taken as bound protein. (**B**) Interaction of LC3A, LC3B, LC3C and GABARAPL2 (GBRL2) with LUVs containing CL. LUVs were composed of ePC:DOPE:CL (33:33:33 mol ratio) + 0.05% Rho-PE. The presence of vesicles and proteins in the different fractions was probed by Rho-PE fluorescence emission and by SDS-PAGE/Coomassie Blue staining respectively. Bars at the bottom: Rho-PE emission was detected only in fractions 3-4 (i.e., floating fractions). The bars at the right-hand side correspond to the percentage of bound protein, taken as protein co-floating with vesicles and calculated by gel densitometry. Data are means ± SD (n ≥ 9). ***P < 0.001

In fact, the *in vitro* interaction of LC3A and LC3C with CL was higher than that of LC3B. Almost all of the LC3C was bound to membranes when CL was present, and, while not all LC3A interacted with CL, it nearly doubled the proportion of bound LC3B (**Fig. 2B**). Binding of the three proteins was dependent both on vesicle curvature (a higher binding was observed to the more highly curved SUV than to LUV) and on CL concentration in the bilayers (**Fig. S1**). Consistent with previous results [**22**] GABARAPL2 did not reach the binding level of the LC3 proteins to CL-containing membranes (**Fig. 2B**). The results indicated that LC3B was not the only member of the LC3 subfamily able to bind CL in lipid bilayers, thus perhaps LC3B might not be the only one able to recognize the externalized CL in damaged mitochondria.

### LC3A and LC3B interact preferentially with CL over other negatively charged phospholipids

To determine whether LC3 interaction with membranes was specific for CL, the binding of these proteins to LUV of different lipid compositions was investigated. None of the LC3 proteins were able to float in the absence of vesicles, or in the presence of liposomes composed of electrically neutral phospholipids [phosphatidylcholine (ePC) and phosphatidylethanolamine (DOPE)] (**Fig. 3A**). Moreover, the interaction of LC3 with membranes containing phosphatidic acid (PA) or phosphatidylglycerol (PG), phospholipids that are structurally and metabolically related to CL, was assayed. PA and PG are also negatively charged, but they contain ≈1 charge per molecule under physiological conditions, vs. ≈2 in the case of CL [**28**]. Neither LC3A nor LC3B did interact with PA- or PG-containing membranes. LC3C was able to bind PA-containing bilayers, albeit the interaction was lower than with CL-vesicles. PG did not increase LC3C binding to liposomes (**Fig. 3**).

In addition, the interaction with phosphatidylinositol 4-phosphate (PI4P) was measured. PI4P contains two negative charges per molecule, as in the case of CL, but two fatty acyl chains per molecule, vs. four in the case of CL. None of the LC3 subfamily members showed the ability to interact with PI4P-containing membranes, suggesting that the interaction could not be explained by purely electrostatic forces (**Fig. 3**). Note that PI4P generation in autophagosomes was shown to be critically important for GABARAP-mediated fusion with lysosomes [**29**], another example of differentiation in the LC3/GABARAP family of proteins. In summary, the above results suggest that all three LC3 subfamily members interact preferentially with CL over other negatively-charged lipids, although LC3C displays also a rather high affinity towards PA, which suggests that this protein is not as specific for CL as the other LC3 isoforms.

**Figure 3.**
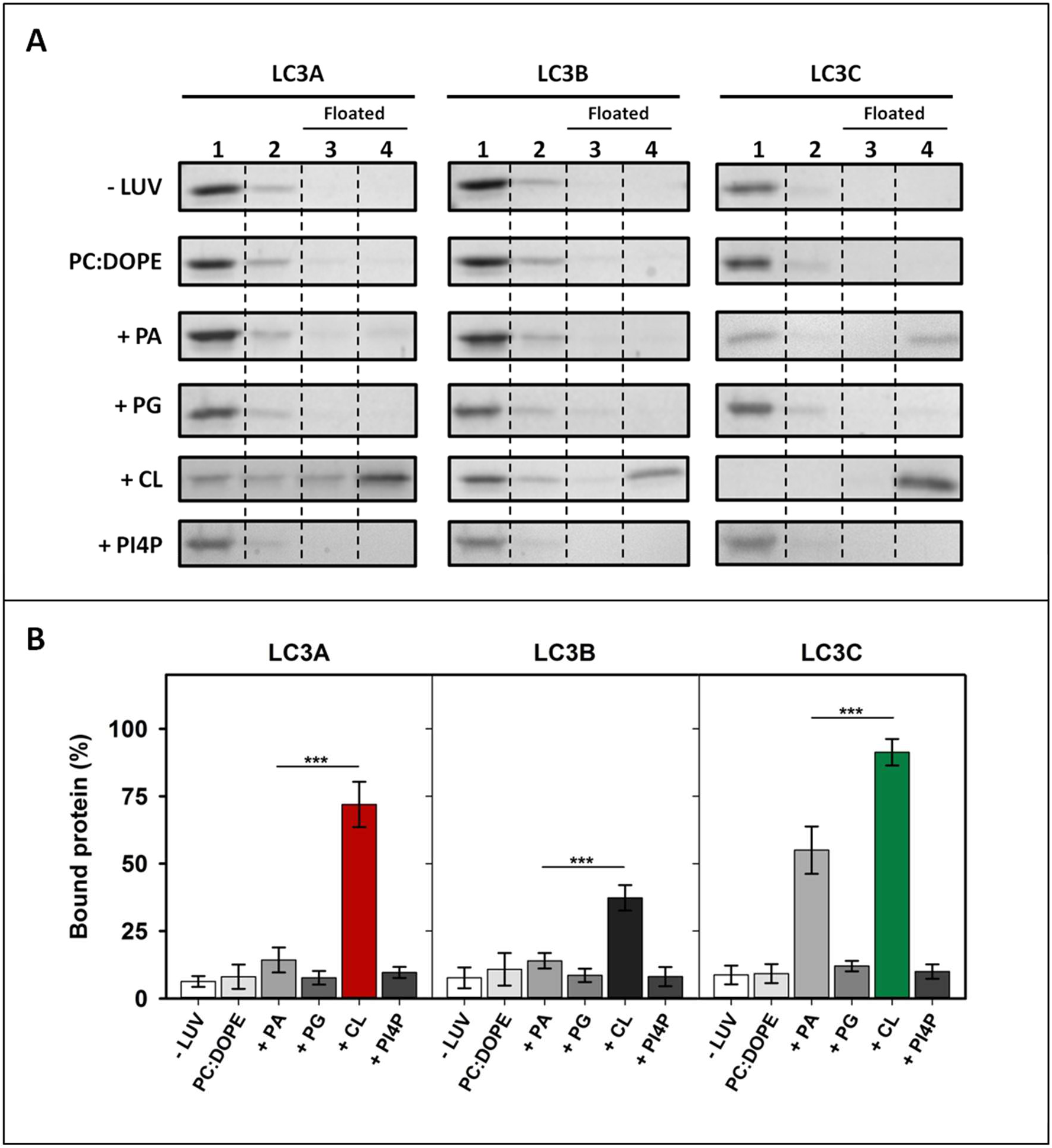
LC3A and LC3B exhibit a marked specificity for CL. Interaction of LC3 proteins with membranes of different compositions, measured by a vesicle flotation assay. (**A**) Representative SDS-PAGE/Coomassie Blue stained gels of the fractions obtained from LC3A, LC3B or LC3C vesicle flotation assays performed without liposomes (-LUVs) or with liposomes of different compositions, either ePC:DOPE (50:50) or ePC:DOPE:X (33:33:33 mol ratio) where X was PA, PG, CL or PI4P. Bound protein was computed as the proportion retrieved in fractions 3+4 (See Fig. 2A). (**B**) Binding of LC3A, LC3B, and LC3C to liposomes quantified by gel densitometry. Data are means ± SD (n ≥ 3). ***P < 0.001.

### The high capacity of LC3C to interact with CL resides in its N-terminal region

To determine the characteristics that made LC3C interact with CL with a higher affinity than LC3B, a sequence alignment of LC3C and LC3B was performed. LC3C exhibited large differences with LC3B in its amino acid sequence (46/72% identity/similarity) (**Fig. 4A**). The main differences were found in the N-terminal region. This observation is consistent with the previously mentioned role of the N-terminal region in the interaction with CL [**16,22**]. Therefore, the possibility that this region could participate in the interaction with CL was tested. For this purpose, a chimaera protein was constructed, containing the LC3C N-terminal region (1-33) and the LC3B 29-120 residues (**Fig. 4B**). The proportion of this chimaera bound to CL-containing liposomes was larger than that of LC3B, reaching the level of LC3C (**Fig. 4C, D**), thus meaning that the N-terminal region of LC3C alone was enough to explain the higher degree of interaction with CL observed with LC3C.

**Figure 4.**
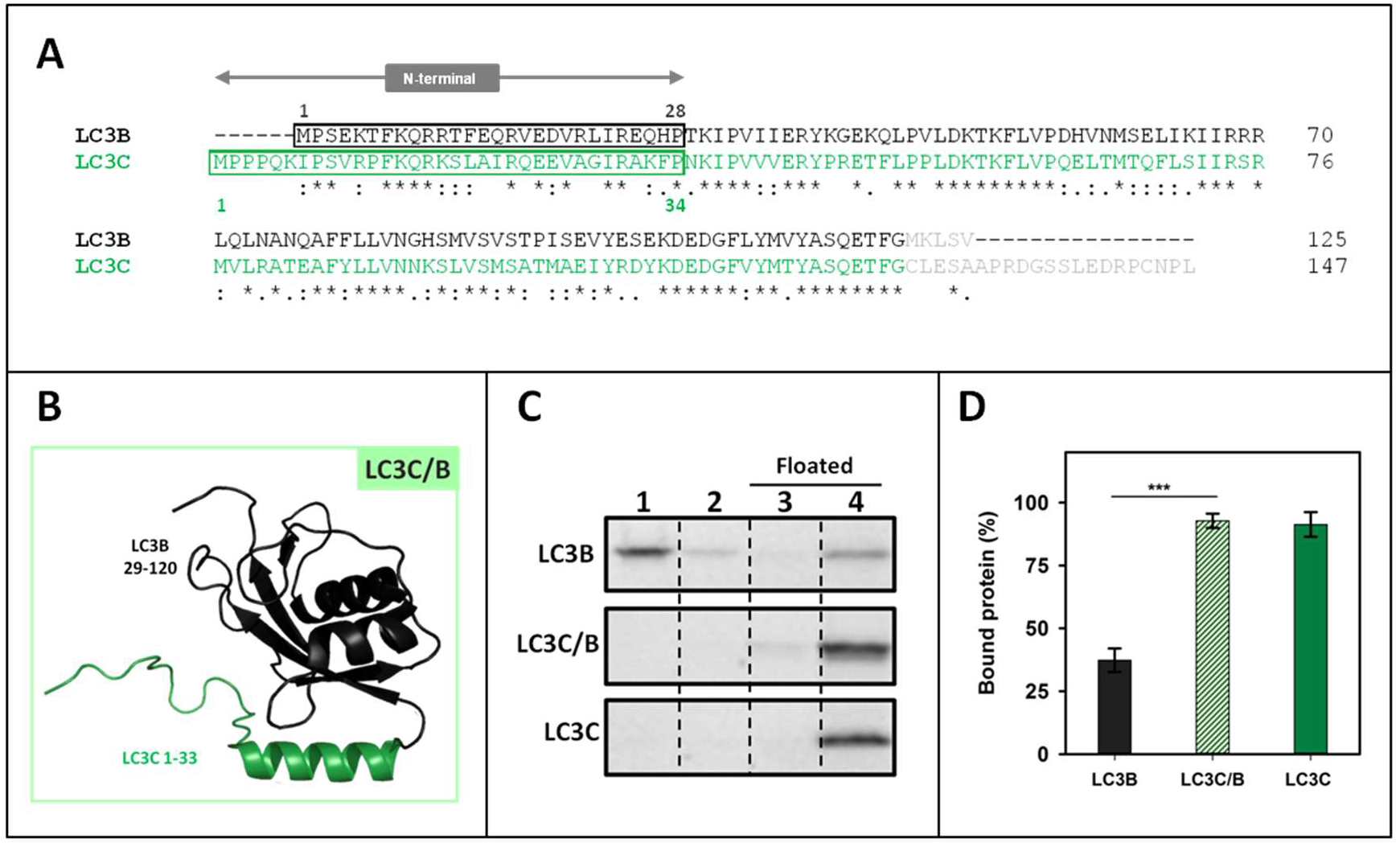
The high capacity of LC3C to interact with CL resides in its N-terminal region. **(A)** Clustal W alignment of LC3B and LC3C amino acid sequences. The N-terminal region of both proteins is boxed. **(B)** 3D outline of the LC3C/B chimaera, composed of the structures of the LC3C N-terminal region (1-33 amino acids) and 29-120 residues of LC3B. **(C)** Representative SDS-PAGE/Coomassie Blue-stained gels of the fractions obtained from LC3B, LC3C/B or LC3C vesicle flotation assays performed with CL-containing liposomes (ePC:DOPE:CL (33:33:33 mol ratio)). Bound protein was computed as the proportion retrieved in fractions 3+4 (See Fig. 2A) (**D**) Binding of LC3B, LC3C/B, and LC3C to CL-containing liposomes quantified by gel densitometry. Data are means ± SD (n ≥ 7). ***P < 0.001.

### The N-terminal regions of LC3A and LC3B are important for their differential interaction with CL

The reason why LC3A interacted with CL with a higher affinity than LC3B was also explored. These two proteins exhibited a 79/92% identity/similarity when their amino acid sequences were compared (**Fig. 5A**) and the main differences were also found in the N-terminal region. Therefore, the implication of this region in LC3A interaction with CL was tested. In this case, a chimaera composed of the LC3A N-terminal region (1-28) and the LC3B protein 29-120 residues was designed (**Fig. 5B**). The chimaera showed a higher interaction with cardiolipin than LC3B, even if it did not reach LC3A binding levels (**Fig. 5C, D**). These results suggested a mode of interaction with CL involving other regions of the protein apart from its N-terminus, similar to the mechanism proposed for LC3B, but not to the one observed with LC3C. The increased CL interaction with the LC3A/B chimaera, as compared to that of LC3B, suggests that LC3A might contain some residues in its N-terminal region allowing a higher interaction with that phospholipid.

**Figure 5.**
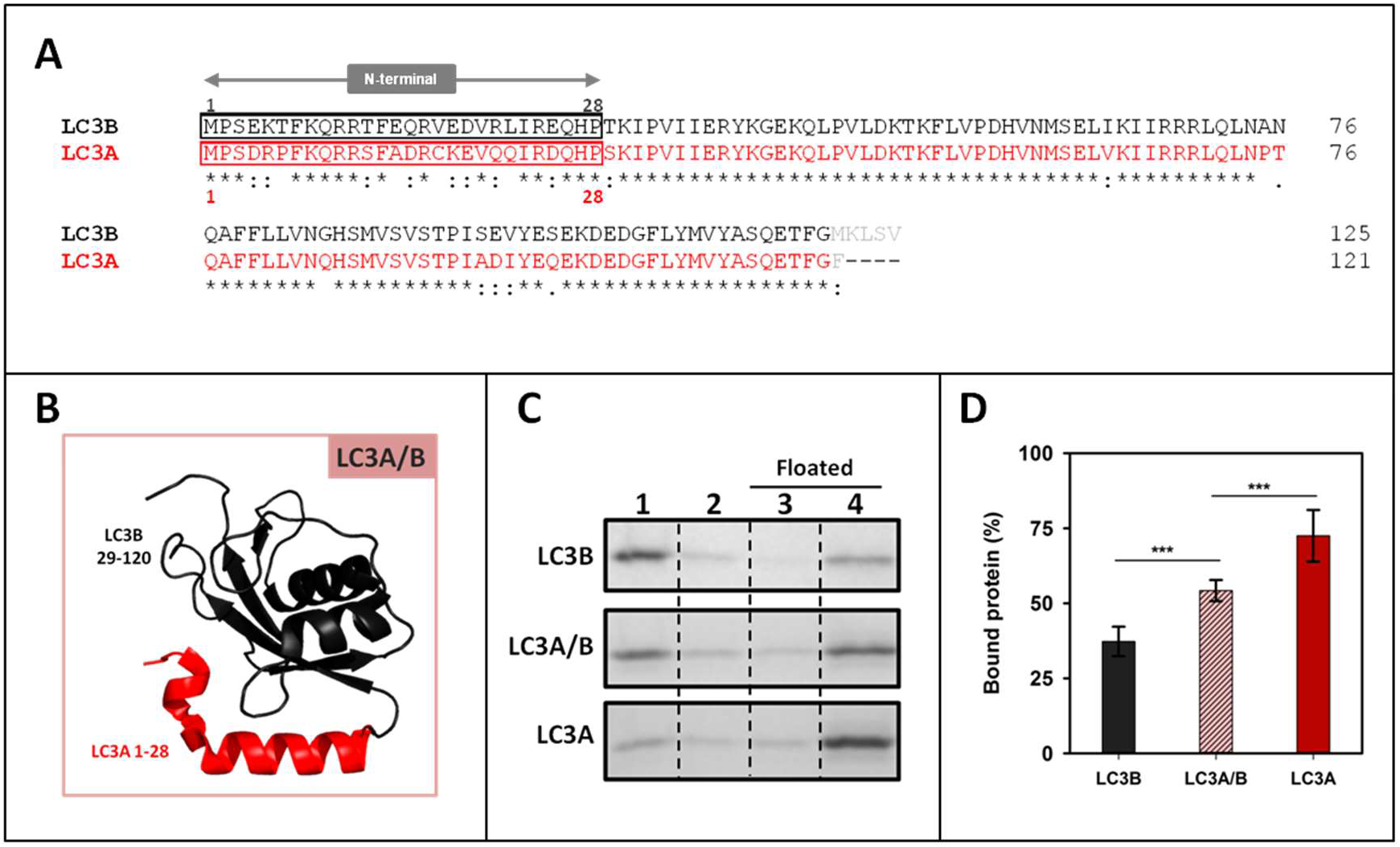
The N-terminal regions of LC3A and LC3B are important for their differential interaction with CL. **(A)** Clustal W alignment of LC3B and LC3A amino acid sequences. The N-terminal region of both proteins is boxed. **(B)** 3D outline of the LC3A/B chimaera, composed of the structures of the LC3A N-terminal region (1-28 amino acids) and 29-120 residues of LC3B. **(C)** Representative SDS-PAGE/Coomassie Blue-stained gels of the fractions obtained from LC3B, LC3A/B or LC3A vesicle flotation assays performed with CL-containing liposomes (ePC:DOPE:CL (33:33:33 mol ratio)). Bound protein was computed as the proportion retrieved in 3+4 (See Fig. 2A) (**D**) Binding of LC3B, LC3A/B, and LC3B to CL-containing liposomes quantified by gel densitometry. Data are means ± SD (n ≥ 8) ***P < 0.001.

### A14 and K18 residues in LC3A N-terminal region are key for its higher interaction with CL

To further analyse the importance of specific residues in the N-terminus of LC3A, a sequence alignment focusing on this region of LC3B and LC3A proteins (**Fig. 6A**) was performed. This comparison revealed that residues predicted by Chu et *al* [**16**] to be involved in the interaction with CL were the same in both proteins (R10, R11), suggesting that the cause for the different behaviour had to be found elsewhere. Two negatively charged amino acids were observed in LC3B (E14, E18) that were not present in LC3A, the latter protein containing instead one neutral and one positively charged residue (A14, K18) (**Fig. 6A, B**). This could be significant considering that the object of the study was an interaction with a negatively charged phospholipid. A more detailed investigation of the effect of ionic strength on LC3-protein binding to CL-containing bilayers was carried out using either the standard buffer (150 mM NaCl) or a buffer containing 300 mM NaCl. The results in **Fig. S2** indicated in all cases a marked decrease of LC3 binding in the presence of high ionic-strength buffer, thus pointing to a role for electrostatic interactions in LC3 binding to CL-containing bilayers.

**Figure 6.**
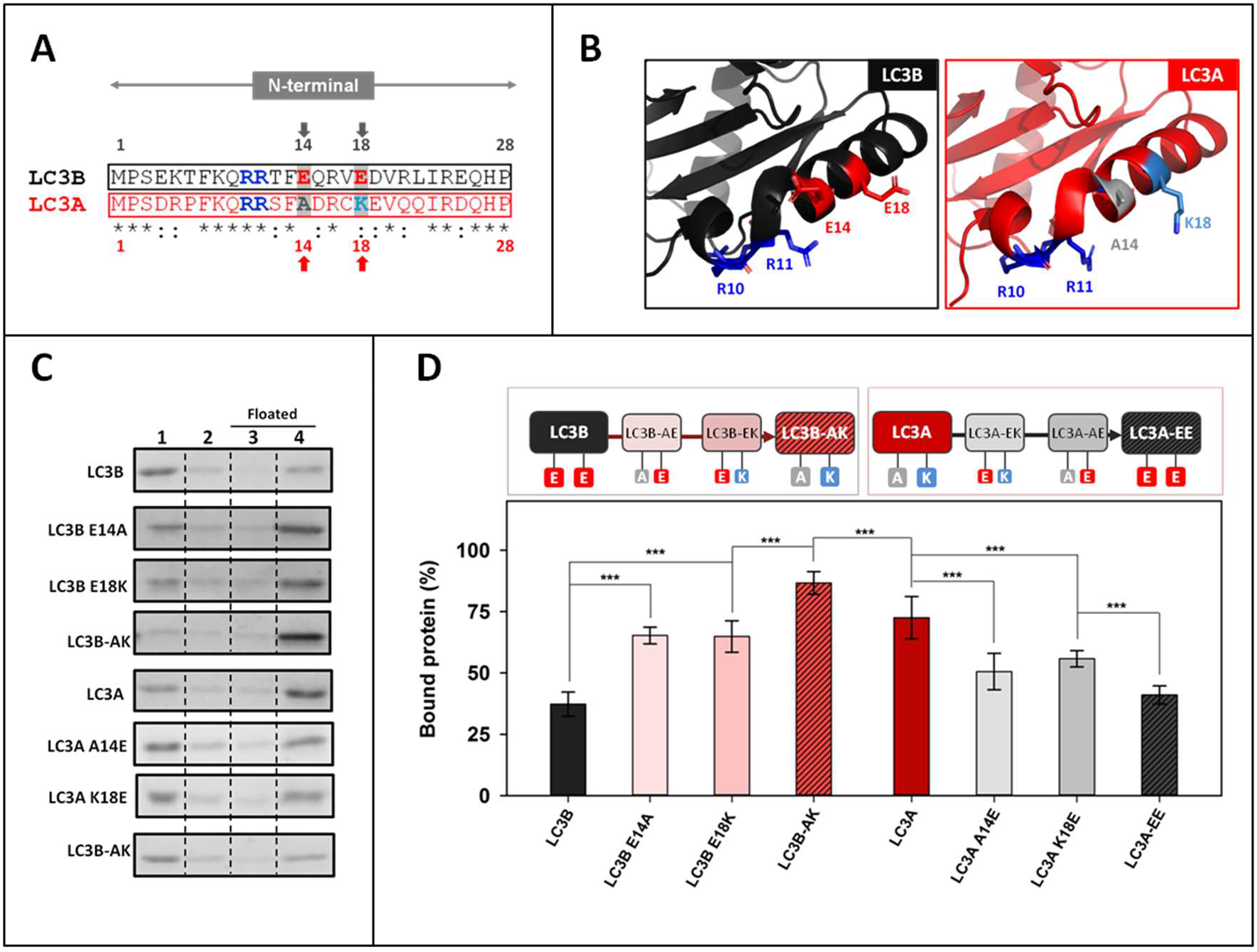
A14 and K18 residues in LC3A N-terminal region are key for its high interaction with CL. (**A**) Comparative analysis of the N-terminal regions of LC3B and LC3A obtained using Clustal W. Amino acids previously proposed to be important in LC3B-CL interaction (R10, R11) and the ones chosen in this study to be mutated (positions 14 and 18) are written in bold and colored; red: negatively charged, blue: positively charged, grey: no charge **(B)** 3D structures of the N-terminal regions of LC3B and LC3A showing the amino acids chosen for this study and the residues already proposed to be involved in the interaction with CL. **(C)** Representative SDS-PAGE/Coomassie Blue-stained gels of the fractions obtained from LC3B, LC3B-AE (LC3B E14A), LC3B-EK (LC3B E18K), LC3B-AK (LC3B E14A E18K), LC3A, LC3A-EK (LC3A A14E), LC3A-AE (LC3A K18E), LC3A-EE (LC3A A14E K18E) vesicle flotation assays performed with CL-containing liposomes (ePC:DOPE:CL (33:33:33 mol ratio)). Bound protein was computed as the proportion retrieved in fractions 3+4 (see Fig. 2A). **(D)** Binding percentage of LC3B, LC3B E14A, LC3B E18K, LC3B-AK (LC3B E14A E18K), LC3A, LC3A A14E, LC3A K18E, LC3A-EE (LC3A A14E K18E) to CL-containing liposomes quantified by gel densitometry. Data shown as mean ± SD (n ≥ 5) ANOVA statistical analysis, ***P < 0.001

To test the involvement of residues 14 and 18 in the differential LC3A and LC3B binding to CL, a series of mutants was designed, expressed and purified. The objective was to “transform” LC3B into LC3A and *vice versa* LC3A into LC3B by mutating the mentioned residues, to elucidate whether the CL-binding affinity was influenced or not. As shown in **Fig. 6C** and **D**, after changing E14 of LC3B into the corresponding A14 of LC3A, i.e. obtaining the mutant LC3B-AE (LC3B E14A), the percentage of bound protein was higher than with native LC3B. Mutating residue 18, to obtain LC3B-EK (LC3B E18K), also increased binding, to the same extent than for LC3B-AE. The highest proportion of bound protein was found with LC3B-AK (LC3B E14A E18K), above the value observed for LC3A (**Fig. 6C, D**).

In the case of LC3A, both single-residue mutants [LC3A-EK (A14E), LC3A-AE (K18E)] exhibited a decreased interaction. Furthermore, the LC3A-EE (LC3A A14E K18E) showed a degree of binding to CL-containing vesicles similar to that of LC3B (**Fig. 6C, D**). Thus, while mutating the two residues of LC3B almost doubled its interaction, changing the two amino acids of LC3A significantly decreased its interaction with CL, demonstrating the importance of the nature of residues 14 and 18 in the higher CL binding activity of LC3A. In addition, when the equivalent of the LC3B E14 residue (A20) in the N-terminal region of LC3C was mutated, to obtain LC3C A20E (**Fig. S3 A, B**), LC3C binding was not affected (**Fig. S3C, D**). Thus, the very effective binding of LC3C to CL-containing vesicles should not be interpreted in the same terms described for LC3A and LC3B above, rather indicating that the whole N-terminal region of LC3C was important for its interaction with CL.

### LC3A and LC3B puncta per cell and their co-localization with mitochondria increase with rotenone treatment

To assess whether the above *in vitro* results were indicative of the behaviour of the proteins in cells, the involvement of each LC3 member in CL-mediated mitophagy was tested (**Fig. 7**). The mechanism controlling CL externalization is not completely understood. CL is a phospholipid mainly found in the inner layer of IMM, therefore 3 steps are required for it to be fully translocated to the external layer of the OMM. Translocation between layers is believed to be catalyzed by phospholipid scramblase 3 (PLS3) [**16,30**]. The transfer between the IMM outer monolayer and the inner leaflet of OMM is thought to be carried out by kinase NDPK-D (also called Nm23-H4) [**31,32**]. It was shown that this externalization was induced by rotenone and 6-OHDA treatments, which did not activate the PINK1-parkin pathway. Moreover, externalization also occurred with carbonyl cyanide m-chlorophenyl hydrazone (CCCP) treatments that depolarized mitochondria and did activate the parkin-dependent pathway [**31**]. These results would point to CL-externalization as an enhancer to attract the autophagic machinery into different mitophagy pathways. To promote a marked CL externalization to the OMM, a treatment with non-lethal doses of rotenone was used. Experiments were performed using LC3A, LC3B, LC3C, and GABARAPL2 to test whether this treatment induced an increase in LC3/GABARAP puncta inside the cell (as a signal of autophagy) and whether it increased the number of puncta that colocalized with mitochondria (mitophagy). The experimental conditions were the same as the ones described in previous studies where the externalization of CL had been shown [**16,31**]. Cells were transfected with LC3 GFP-tagged proteins and treated with 1 μM rotenone. GFP-GABARAPL2 was used as a control (this protein is unable to translocate to mitochondria upon rotenone treatment in U87MG cells) [**22**].

**Figure 7.**
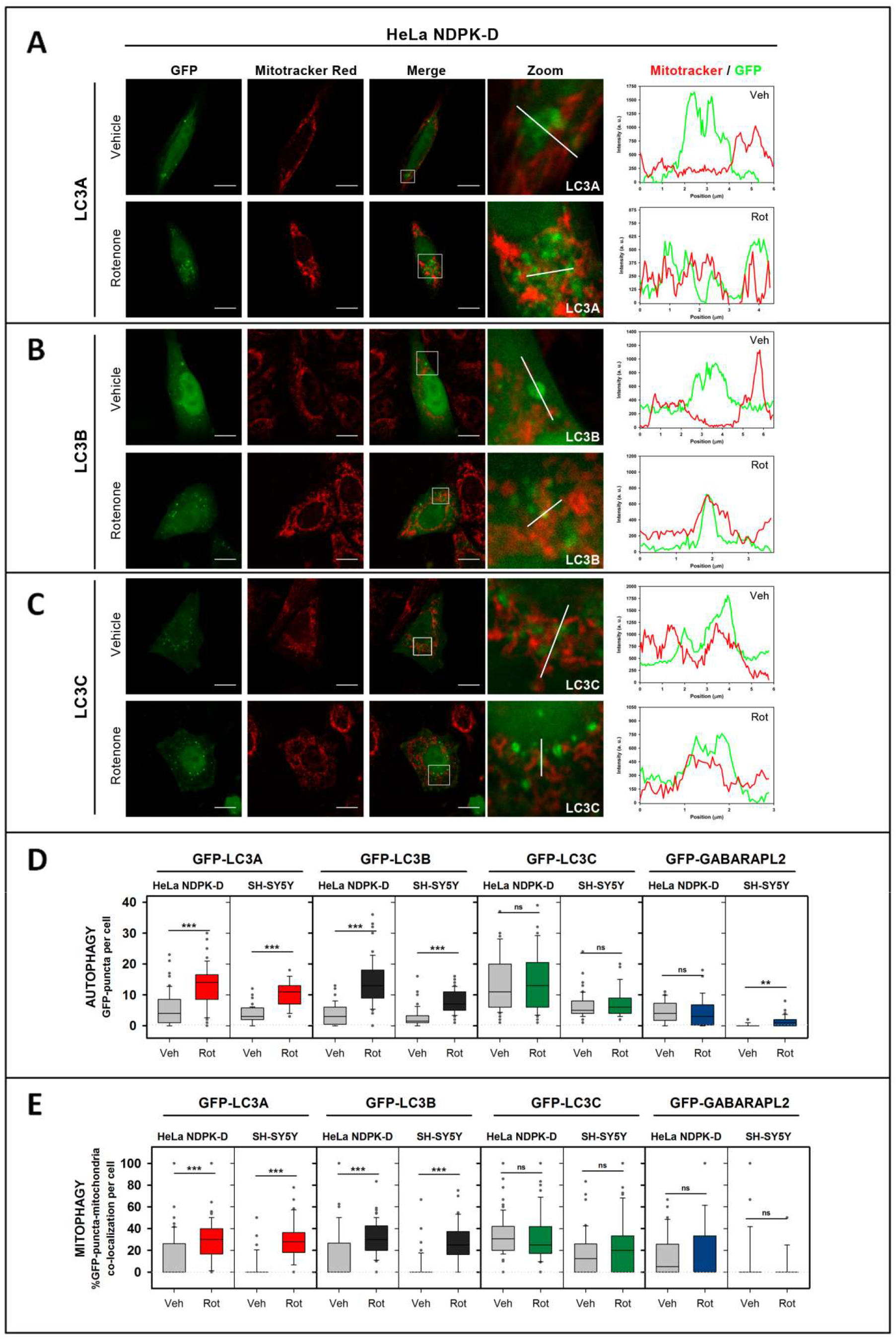
LC3A and LC3B puncta per cell and their co-localization with mitochondria increase with rotenone treatment. Cells were transfected with different members of the LC3/GABARAP family tagged with GFP. Mitochondria were labeled using Mitotracker Red, prior to treatment with 1 μM rotenone. Vehicle (Veh) controls were treated with DMSO. Images were acquired using a Nikon Eclipse C1 confocal microscope. Scale bar: 10 μm. At the right-hand side of each condition, Mitotracker (red) and GFP (green) line profiles show examples of colocalization and non-colocalization events. (**A-C**) Representative images of GFP-LC3 HeLa NDPK-D transfected cells, untreated (vehicle) and treated with rotenone. (**D**) Numbers of GFP puncta per cell, an indication of autophagy, in HeLa NDPK-D and SH-SY5Y cells (representative images of the latter in Fig. S5). (**E**) Percent GFP puncta that colocalize with mitochondria, a signal of mitophagy, in HeLa NDPK-D and SH-SY5Y cells (representative images of the latter in Fig. S5). To estimate the percent colocalization, images were analyzed with JACop plugging of Image J. At least 30 images were analyzed per condition. ***P < 0.001, **P < 0.01, ns: non-significant.

HeLa and SH-SY5Y neuroblastoma cells were used to test the LC3 subfamily members role in rotenone-induced mitophagy. The basal expression of one of the proteins involved in CL translocation, NDPK-D, is low in normal HeLa cells, therefore a HeLa cell line stably expressing this protein was used [**31**]. Results using HeLa NDPK-D cell line showed that GFP-LC3A participates in rotenone-induced autophagy and mitophagy (**Fig. 7A, D, E**), consistent with its ability to interact with CL-containing membranes. Moreover, in agreement with the previously published results [**16,22**], although rotenone treatment increased GFP-LC3B puncta and colocalization with mitochondria (**Fig. 7B, D, E**), it had no effect on GFP-GABARAPL2 (**Fig. 7D, E, S4A**). A high amount of GFP-LC3C puncta was observed under basal conditions (**Fig. 7C, D, E**), and despite the remarkable level of interaction displayed by LC3C with CL-containing membranes (**Fig. 2B**), the amount of GFP-LC3C puncta did not increase after rotenone treatment (**Fig. 7C, D, E**). As seen in **Fig. 7D** and **E**, equivalent assays performed using SH-SY5Y cells showed similar results with all LC3 proteins. In the case of GFP-GL2 in SH-SY5Y cells, rotenone causes a small significant increase in puncta/cell, but it did not correspond with a significant increase in colocalization (**Fig. 7D, E, S4B**). The similar results obtained using two different cell lines suggest that LC3A and LC3B are able to recognize CL not only in model membranes but also in damaged mitochondria, and that LC3A and LC3B, but not LC3C or GABARAPL2, are involved in rotenone-activated mitophagy.

The formation of puncta, detected by conventional fluorescence microscopy, has been interpreted in the above experiments as an indicator of soluble LC3 (LC3-I) conversion to the lipidated and autophagosome-associated form (LC3-II). To ascertain that the apparent autophagy observed was indeed due to autophagy vesicles and not to aggregates, non-conjugatable mutants of the proteins were used. In these mutants the C-terminal glycine had been mutated into an alanine residue, resulting in a non-functional form of the proteins that could not be conjugated to PE, thus it could not be anchored to the autophagosomal membrane. Consistently, the mutant proteins failed to give rise to puncta, indicating that the GFP signals detected in **Fig. 7** did not correspond to nonspecific aggregates (**Fig. S5**).

### The double mutation that hampers LC3A binding to CL *in vitro* also decreases its location to mitochondria in mitophagy

To determine whether the residues identified as key for the differential *in vitro* interaction of LC3A and LC3B with CL were also important in cells, the ability of the double mutants to recognize damaged mitochondria after rotenone treatment was tested. NDPK-D-expressing HeLa cells were transfected with the GFP-tagged LC3 WT and mutant proteins (LC3A, LC3A-EE, LC3B, and LC3B-AK) and treated with rotenone. Consistent with the *in vitro* results, in which LC3A-EE exhibited much lower binding to CL vesicles than its native counterpart (**Fig. 6C, D**), cells transfected with GFP-LC3A-EE showed lower colocalization levels with mitochondria (**Fig. 8A, C, S6D**). The substitution of those two amino acids in LC3B (LC3B-AK), that exerted a significant positive action *in vitro* (**Fig. 6C, D**), did not have a parallel effect in cells (**Fig. 8B, D, S6D**). Similar experiments, with comparable results, were performed with SH-SY5Y cells (**Fig. S6A, B, C**). The overall results indicated that the A14 and K18 residues of LC3A, that caused the difference between LC3A and LC3B in the *in vitro* interaction with CL, were also important for LC3A interaction with externalized CL in cells.

**Figure 8.**
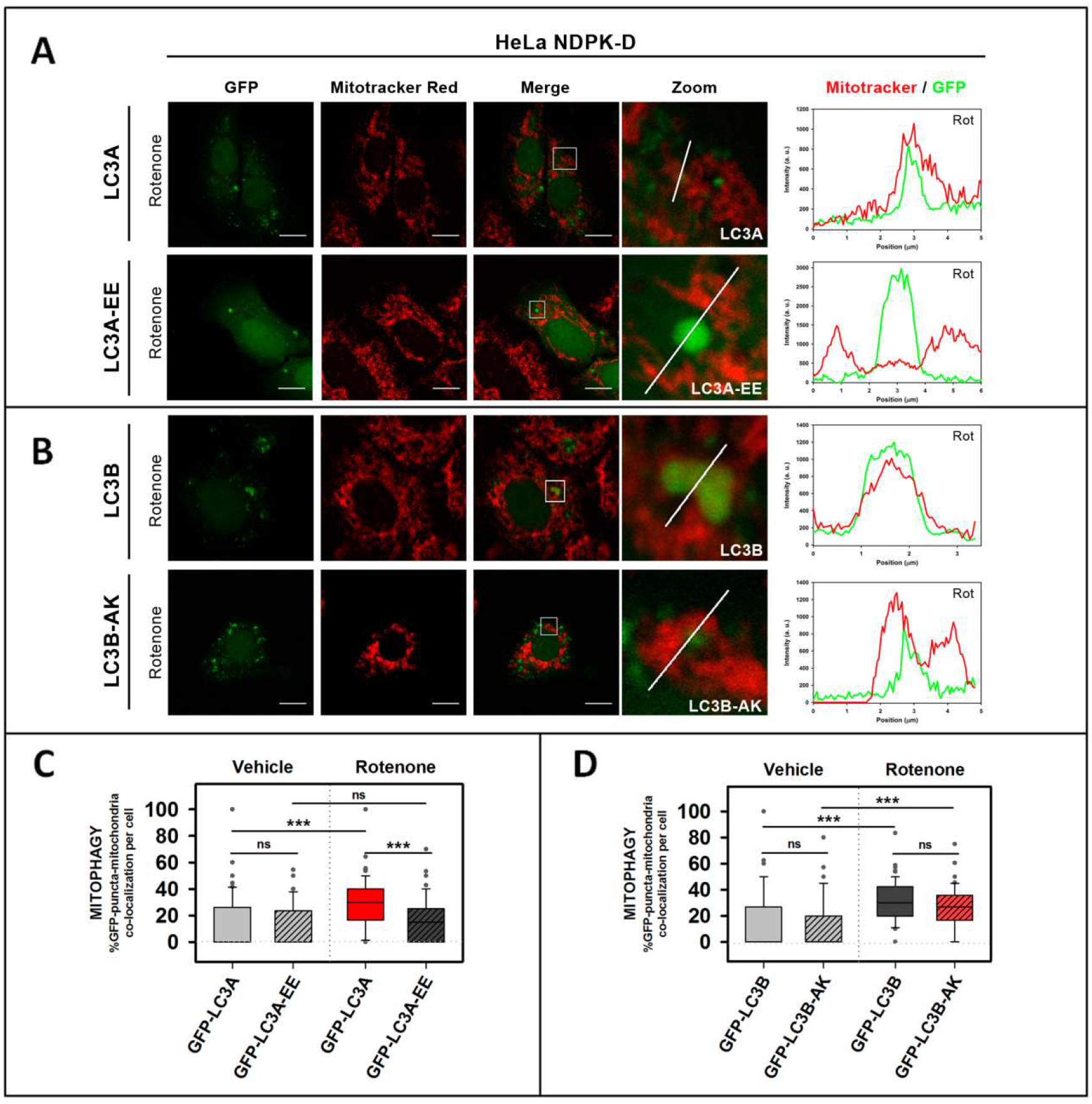
The LC3A-EE double mutation that hampers LC3A binding to CL *in vitro* also decreases its location to mitochondria in mitophagy. NDPK-D-expressing HeLa cells were transfected with GFP-tagged WT or mutant LC3. Mitochondria were labeled using Mitotracker Red, prior to treatment with 1 μM rotenone. Vehicle (Veh) controls were treated with DMSO. Images were acquired using a Nikon Eclipse C1 confocal microscope. Scale bar: 10 μm. At the right-hand side of each condition, Mitotracker (red) and GFP (green) line profiles show examples of colocalization and non-colocalization events. (**A**) Representative images of GFP-LC3A and GFP-LC3A-EE HeLa NDPK-D transfected cells treated with rotenone. (**B**) Representative images of GFP-LC3B and GFP-LC3B-AK HeLa NDPK-D transfected cells treated with rotenone. Representative images of the untreated conditions are in Fig. S6D. (**C-D**) Percent GFP puncta colocalizing with mitochondria in each condition, an indication of mitophagy. To compute the percent colocalization, images were analyzed with JACop plugging of Image J. At least 30 images were analyzed per condition. ***P < 0.001, ns: non-significant.

### LC3A but not LC3B is able to bind oxidized CL

It has been proposed that when CL participates in apoptosis, it is previously oxidized due to the peroxidase activity of cytochrome C [**33**], while the CL fraction that is recognized by the autophagy machinery is not [**16**]. Taking into account that the interplay between autophagy and apoptosis implies a high level of complexity [**34**], understanding how the role of CL is affected by its oxidation state could be helpful in order to understand how these two important processes are regulated.

In order to assess the ability of LC3A and LC3B in recognizing oxidized CL, LUVs were treated with CuCl_2_ for 5 h, and the increase in A245 was measured to quantify lipid oxidation (**Fig. 9A**). The highly unsaturated CL is the main lipid oxidized under those conditions. Then, protein bound to control and oxidized vesicles was measured. The results obtained with LC3B and LC3A in these experiments showed that the binding of LC3B decreased with CL oxidation, but not that of LC3A (**Fig. 9B**). These results suggested that LC3A could recognize oxidized CL in the OMM, perhaps preventing its recognition by the apoptotic machinery.

**Figure 9.**
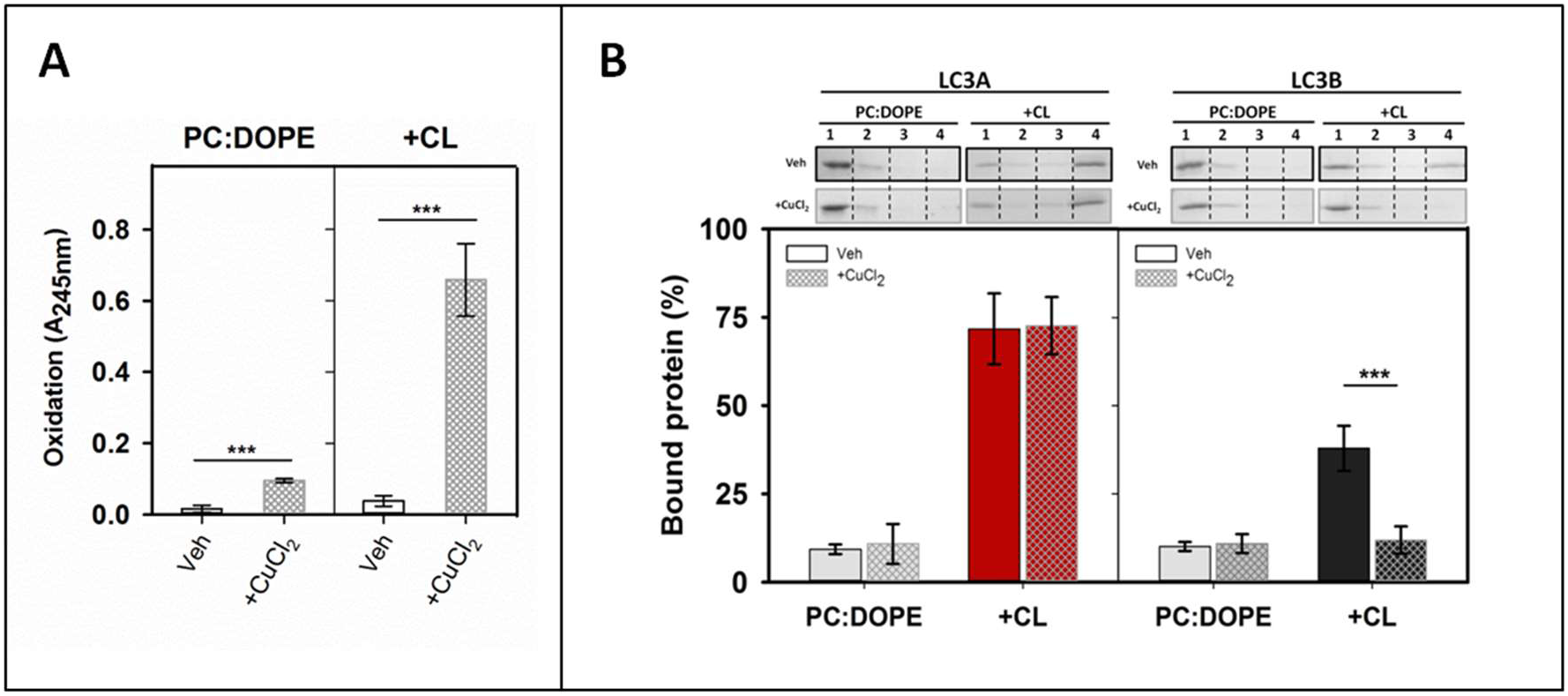
**LC3A, but not LC3B, binds oxidized CL** Interaction of LC3A or LC3B proteins with oxidized membranes measured by a vesicle flotation assay. (**A**) Lipid oxidation was assessed measuring absorbance at 245 nm (A_245nm_) of the ePC:DOPE (50:50) or ePC:DOPE:CL (33:33:33) liposomes, treated without (Veh) or with CuCl_2_ (+CuCb). Data shown as means ± SD (n ≥ 3). ***P < 0.001 (**B**) Top: Representative SDS-PAGE/Coomassie Blue-stained gels of the fractions obtained from LC3A or LC3B vesicle flotation assays performed with liposomes ePC:DOPE (50:50) or ePC:DOPE:CL (33:33:33) which had been previously treated without (Veh) or with CuCl_2_. Bound protein is computed as the proportion retrieved in fractions 3+4 (See Fig. 2A). Bottom: Binding of LC3A, LC3B, and LC3C to non-oxidized and oxidized liposomes, quantified by gel densitometry. Data shown are means ± SD (n ≥ 3). ***P < 0.001.

## Discussion

Deciphering the different modes of signalling damaged mitochondria and the role that each of the LC3/GABARAP family members plays in their recognition is extremely important in understanding mitophagy. Many types of receptors and adaptors participate in this process, and different studies reinforce the idea that the LC3/GABARAP members could have specific functions depending on the particular mechanism involved [**14**]. Experiments performed using LC3/GABARAP family members in hexa- and triple-knockout cells suggest that GABARAP is the main subfamily involved in PINK1-parkin-dependent mitophagy [**35**]. There are examples of proteins that act as mitophagy receptors by interacting preferentially with one or several of the LC3/GABARAP family members, for example, NIX recruits GABARAP [**36**] and GABARAPL1 [**37**], while FUNDC1, BINP3 and Bcl2-L-13 are found bound to LC3B [**38–40**], and FKBP8 is recognized by LC3A [**41**]. In the case of the phospholipid CL acting as a mitophagy receptor, LC3B was shown to be a mediator [**16**], and our previous work comparing both subfamilies suggested that neither GABARAP nor GABARAPL2 were involved [**22**]. However, a comparative study of the different LC3 subfamily members function(s) does not appear to be available.

### The specific interaction of LC3A with CL allows its participation in the mechanism of CL-mediated mitophagy

The interaction of LC3A with membranes was specific for the presence of CL (**Fig. 3**). LC3A did not show any measurable interaction with other lipids containing the same negative charge of CL, such as PI4P (**Fig. 3B**), therefore, this interaction did not seem to be exclusively dependent on the lipid charge. The amino acid sequence of LC3A and LC3B is very similar, and even some residues proposed to be key for the interaction with CL, R10 and R11 [**16**], located in the alpha helix 1, are identical in both proteins (**Fig. 1B, 6A, B**). However, in the above studies with model membranes a clear difference in CL binding was observed between LC3A and LC3B, the former being more active (**Fig. 2B**). Experiments involving the LC3A/B chimaera (**Fig. 5**) and higher salt concentrations (**Fig. S2**), suggested that the higher binding detected with LC3A could be in part due to the existence of positively charged residues in its N-terminal region, that are not present in LC3B. Comparative binding assays using LC3A and LC3B single and double mutants showed that the key amino acids were located in positions 14 and 18 of helix 2 (**Fig. 6**). Therefore, the CL binding site of LC3A appears to be composed of two alpha-helices containing positively charged residues, in agreement with previously described binding sites for other soluble proteins that can also bind CL in membranes [**42**]. However, the fact that LC3A/B chimera did not reach the same binding levels as LC3A (**Fig. 5**) suggests the implication of other protein regions apart from the N-terminus, as previously proposed for LC3B, in the interaction of LC3A with CL.

To further dissect the differences among these subfamily members, the participation of each LC3 subfamily member in CL-mediated mitophagy was studied by assessing the colocalization of transfected GFP-tagged proteins with mitochondria in two different cell lines, HeLa-NDPK-D and SH-SY5Y. Even though the effect of the GFP-tag in the LC3/GABARAP protein function had been criticized [**35**], we decided to use this method because the existing antibodies against LC3 subfamily members were not specific enough. The similar results obtained using anti-LC3 antibodies [**20,22**] and GFP-tagged LC3B [**16,31**] to study CL-mediated mitophagy in cells, supported the notion that GFP-tag did not affect the recognition of externalized CL in damaged mitochondria. In addition, the results obtained with the non-conjugatable mutants of each homolog (**Fig. S5**) in which the number of puncta was residual, showed that the GFP-puncta are, in fact, autophagic vesicles, and not aggregates. The above observations indicate that the results obtained with the cell lines are reliable.

The experiments performed in cells showed that, apart from LC3B, LC3A was also able to recognize mitochondria containing externalized CL (**Fig. 7**), implying that LC3A would recognize damaged mitochondria in a PINK1/parkin-independent manner. The behaviour of LC3A-EE in cells showed that mutations decreasing LC3A-CL binding *in vitro* (**Fig. 6D**), also affected its function in cells, decreasing its ability to colocalize with mitochondria (**Fig. 8C, S6A**). These results indicated that residues 14 and 18 of LC3A were important for damaged mitochondria recognition in cells. The participation of LC3A in parkin-independent mitophagy has been described previously, LC3A being able to specifically recognize the mitophagy receptor FKBP8 located in the OMM of damaged mitochondria [**41**]. However, LC3A recognition of CL and of FKBP8 do not necessarily exclude each other, and these two mechanisms might act together in cooperation. It is possible that CL externalization brings LC3A closer to mitochondria to facilitate binding to FKBP8.

Despite the differences found *in vitro* (**Fig. 2B**), the same behavior was observed for LC3A or LC3B when colocalization with mitochondria was considered (**Fig. 7E**). Moreover, while the exchange of residues 14 and 18 of LC3B for the corresponding ones of LC3A in the LC3B-AK mutant increased significantly its interaction with CL-containing membranes (**Fig. 6D**), this change did not affect its function in mitophagy: the double mutant did not show a higher level of co-localization with mitochondria when compared to LC3B (**Fig. 8D, S6B**). These results could suggest that the mitophagy level observed with LC3B represents a colocalization threshold for the rotenone treatment. If that were the case, even if mitochondrial recognition by LC3A or LC3B-AK were better, it would not lead to an observable higher colocalization. This could also indicate that, at variance with the equivalent residues in LC3A, the negatively charged residues present in the LC3B N-terminal region (E14, E18) do not play a role in the interaction with externalized CL in cells (**Fig. 6B, 8C**), implying that the molecular mechanism by which LC3A and LC3B interact with CL in the cell might be different, or else that their interaction would not be regulated in the same way.

### LC3C does not participate in CL-mediated mitophagy

The interaction of LC3C with membranes was not specific for the presence of CL, as this protein could also interact with PA (**Fig. 3B**). Moreover, despite the higher interaction displayed with CL-containing membranes when compared with LC3A or LC3B (**Fig. 2B**), LC3C does not take part in CL-mediated mitophagy, at least under our experimental conditions (**Fig. 7**). The peculiar behaviour of LC3C is not surprising, it has been shown that LC3C has different functions and targets as compared to the other subfamily members [**4**], such as its specific function in antibacterial autophagy [**43**].

While LC3A and LC3B show a high similarity in their sequence (92%), LC3C is the most differentiated homolog (71-72%). Unlike in other LC3/GABARAP members, the N-terminal region of LC3C does not form a stable α1-helix but a “sticky arm” (**Fig. 1A**) consisting of a polyproline motif [**44**]. Another important difference is that although LC3A and LC3B are ubiquitously expressed in almost all tissues [**45**], LC3C expression is extremely low and/or tissue-specific (lungs and placenta) [**24**]. It has also been shown that the interactome of LC3C is the one that differs the most among the LC3 subfamily [**46**]. The results in this paper show CL-mediated mitophagy as yet another mechanism in which LC3C behaves differently from the other subfamily members.

The assays using HeLa-NDPK-D and SH-SY5Y cells showed that in the basal state GFP-LC3C was already forming puncta (**Fig. 7D**) and some of these puncta colocalized with mitochondria (**Fig. 7E**). Moreover, CL externalization by rotenone did not have any further effect (**Fig. 7E**). These unchanged levels of autophagy and mitophagy after rotenone treatment suggest that CL externalization had little or no effect on the participation of LC3C in autophagy or on its mitochondrial localization. These results are in accordance with a previous study that used Saos-2 cells and GFP-LC3C [**45**]. In the basal state, they also observed more puncta per cell than with LC3A and LC3B, but these values did not increase after starvation-induced autophagy.

The observed GFP-LC3C colocalization with mitochondria could be explained by the recently discovered involvement of LC3C in the maintenance of basal mitochondrial network homeostasis [**47**]. In this study, Le Guerroue *et al*. (2017) [**47**] described a LC3C-driven mitophagy mechanism that mediates autophagosome degradation of mitochondrial parts in a piecemeal way. Moreover, the affinity that LC3C shows towards PA-containing membranes could also allow its interaction with healthy mitochondria, which contain PA in their OMM [**48**]. In addition, previous results indicated that LC3C does not have a relevant function in parkin-mediated mitophagy [**35**]. Those studies, together with the results in this paper suggest that the main function of LC3C in mitophagy could be related to the healthy mitochondrial network maintenance, by regulating degradation of the damaged mitochondrial parts [**47**].

### LC3A could prevent excessive apoptosis activation

It has been shown that CL could interact with various proteins involved in the regulation of crucial cellular and mitochondrial processes [**42**]. Besides LC3A and LC3B interaction with CL studied here, this phospholipid can also interact with Beclin-1 [**49**], a key component in the autophagic core machinery, with Drp1, one of the proteins in charge of mitochondrial fission [**27**], or with OPA-1, involved in IMM fusion [**50**]. Interestingly, CL externalization also occurs during apoptosis [**33**]. Externalized CL can interact with many proteins involved in this process, such as BAK, BAX, or Caspase-8 [**26,51,52**] and it also participates in other programmed cellular death pathways such as NLRP3 inflammasome activation [**53**].

All these studies show that both CL and its location are key for the correct functioning of mitochondria. Among its different roles, it is surprising that CL externalization could promote both a generally “pro-survival” pathway, such as autophagy, and a “pro-death” pathway, such as apoptosis [**54**]. However, it has been proposed that CL participating in apoptosis is previously oxidized due to the peroxidase activity of cytochrome C [**33**], while CL recognized by the autophagy machinery is not [**16**]. Interestingly, the results obtained with LC3B and LC3A in the experiments with oxidized CL show that while LC3B binding decreases when CL is oxidized, CL oxidation does not affect LC3A binding (**Fig. 9**). This result might indicate that LC3A could recognize oxidized CL in the OMM, shielding it from the apoptotic machinery. Therefore, LC3A might act as a counterweight, preventing excessive activation of apoptosis. According to our *in vitro* data, LC3B would not be able to achieve the latter function, as its ability to interact with CL decreases with oxidation.

### Concluding remarks

The importance of CL externalization as a signal to degrade damaged mitochondria via autophagy has been demonstrated *in vitro* and *in vivo* [**16,18,22,31**]. In this work, LC3A has been identified as an additional LC3 family member involved in mitophagy, and key residues for its interaction with CL have been singled out. This contribution could facilitate the design of precise modulators for this mitophagy mechanism. However, the possibility of this process being cell-, tissue- or organ-specific, or even the possibility of a cross-talk between different types of autophagy cannot be dismissed. Further investigations would be required to improve our understanding of the mechanisms triggering mitophagy, which could in turn be involved in the appearance of important neurodegenerative diseases such as Parkinson’s or Alzheimer’s.

## Materials and methods

### Materials

L-α-phosphatidylcholine from hen egg yolk (ePC, 840051), heart bovine cardiolipin (CL; 90% tetralinoleoylcardiolipin, 5% tetraoleoylcardiolipin, 5% unknown; 840012), phosphatidylinositol-4-phosphate (PI4P, 840045), egg phosphatidic acid (PA; 840101), egg L-α-phosphatidylglycerol (PG; 841138), 1,2-dioleoyl-sn-glycero-3-phosphatidylethanolamine-N-lissamine rhodamine B sulfonyl) (Rho-PE, 810150), and 1,2-dioleoyl-*sn*-glycero-3-phosphatidylethanolamine (DOPE, 850725) were purchased from Avanti Polar Lipids, Inc. (Alabaster, AL). DMEM medium, RIPA 1640 medium, Lipofectamine™ 3000 (L3000008), MitoTracker™ Red CMXRos (M7512), Fetal Bovine Serum (10270106), non-essential amino acids (11140050) and penicillinstreptomycin (15140122) were purchased from Thermo-Fisher Scientific (Waltham, MA). Rotenone (ab143145) was purchased from Abcam (Cambridge, UK).

### DNA constructs and site-directed mutagenesis

The pGEX4T-1 plasmids for expression of several of the various Atg 8 orthologs tagged with glutathione S-transferase (GST) (human LC3A, human LC3B, human LC3C and human GABARAPL2) were kindly provided by Dr. Ivanna Novak (School of Medicine, University of Split, Croatia). Note that each of these Atg8 orthologs was a truncated form lacking the C-terminal Gly. The Gly-exposed forms, such that no ATG4-mediated preprocessing was necessary, that were used in this work, were constructed using a QuikChange site-directed mutagenesis kit (Stratagene, San Diego, CA, 200514). The green fluorescent protein (GFP) plasmids (pHAGE-N-eGFP) were kindly provided by Dr. Christian Behrends (Munich Cluster for Systems Neurology, Munich, Germany.) Note that each of these GFP-tagged Atg8 orthologs contains the full-length sequence of the proteins. For the preparation of the mutant proteins LC3A A14E (LC3A-EK), LC3A K18E (LC3A-AE), LC3A A14E K18E (LC3A-EE), LC3B E14A (LC3B-AE), LC3B E18K (LC3B-EK), LC3B E14A E18K (LC3B-AK), LC3C A20E, GFP-LC3A G120A, GFP-LC3B G120A, GFP-LC3C G126A and GFP-GL2 G116C, an inverse PCR-based site-directed mutagenesis using the KOD-Plus mutagenesis kit (Toyobo, Osaka, Japan, SMK-101) was employed, following the instructions of the manufacturer. The sequences in all mutant constructs were confirmed by DNA sequencing analysis (Secugen S.L., Madrid, Spain). The primers used for making the different mutants are listed in **STable 1**. The chimaeras (LC3A/B, LC3C/B) and the GFP-tagged double mutants (GFP-LC3A-EE, GFP-LC3B-AK) were obtained by subcloning (synthesized by GenScript, Piscataway, NJ).

### Recombinant protein expression and purification

All proteins were purified from soluble fractions of bacterial extracts obtained in the absence of detergents, and they were >90% pure as evaluated by Coomassie Blue-stained SDS-PAGE. *E. coli* BL21(λDE3) cells were transformed with appropriate plasmids, they were grown to OD_600_=0.8 and induced with 0.5 mM IPTG for 16 h at 20 °C. Following centrifugation at 4,500 RCF for 15 min, the pellet was resuspended and sonicated in breaking buffer (phosphate buffered saline (PBS) with protease inhibitors mixture and 1mM DTT). After removal of cellular debris by centrifugation at 30,000 RCF for 30 min at 4 °C, the sample supernatant fraction was incubated with 1 ml Glutathione Sepharose 4B (GE Healthcare, 17-0756-01) for 3 h at 4 °C to bind GST-tagged proteins. Bound proteins were cleaved with Thrombin Protease (GE Healthcare, 27-0846-01) overnight at room temperature in Thrombin Buffer (140 mM NaCl, 2.7 mM KCl, 10 mM Na_2_HPO_4_, 1.8 mM KH_2_PO_4_ (pH 7.3) with freshly added 1 mM DTT). After cleavage they were eluted in Assay Buffer (50 mM Tris-HCl pH 7.5, 150 mM NaCl, 1 mM EDTA with freshly added 1 mM DTT), then concentrated to 500 μl using Amicon Ultra-4 (4 mL, 3 kDa cut-off) (Millipore, Darmstadt, Germany, UFC800324) and loaded onto a Superdex75 10/300 GL size exclusion column (GE Healthcare, Buckinghamshire, UK, GE17-5174-01) equilibrated in Assay Buffer supplemented with freshly added 1 mM DTT. Proteins were aliquoted, flash-frozen and stored in 20% glycerol at −80 °C until further use.

### Liposome preparation

The appropriate lipids were mixed in organic solution and the solvent was evaporated to dryness under a N2 stream. Then the sample was kept under vacuum for 1 h to remove solvent traces. The lipids were swollen in Assay Buffer in order to obtain multilamellar vesicles (MLVs). Large unilamellar vesicles (LUV) were produced from MLV according to the extrusion method described by Mayer et al. (1986) [**55**]. They were subjected to 10 freeze/thaw cycles, and then extruded [LIPEX Liposome Extrusion System (Transferra Nanosciences, Burnaby, Canada)] using 0.1-μm pore size Nuclepore filters (Whatman, 110605). Small unilamellar vesicles (SUV) were obtained by sonicating MLV with a probe tip sonicator (MSE Soniprep 150, MSE, UK) for 20 min (10 sec on, 10 sec off) on ice. Vesicle size was checked by quasi-elastic light scattering using a Malvern Zeta-Sizer 4 spectrometer (Malvern Instruments, Malvern, UK). LUV had an average diameter of ≈100 nm and SUV average diameter was ≈50 nm. Phospholipid concentration was determined by phosphate analysis [**56**].

### Vesicle flotation assay

Protein interaction with membranes was assessed using flotation in sucrose gradients (See **Fig. 2A**). Liposomes (3 mM), containing 0.05 mol% Rho-PE for detection, were incubated with 10 μM of the different purified proteins, for 1 h at 37 °C in Assay Buffer under continuous stirring (1100 rpm). The protein/lipid mix was adjusted to a 1.4 M sucrose concentration in a final volume of 300 μl and transferred to a centrifuge tube. This first (bottom) layer was overlaid with successive solutions containing 0.8 M (400 μl) and 0.5 M (300 μl) sucrose. The three-layer gradients were centrifuged in a TLA-120.2 rotor (Beckman Coulter, Brea, CA, US) at 355040 RCF (avg) for 50 min at 4 °C. After centrifugation, four 250-μl fractions were collected starting from the bottom. Proteins were detected in SDS-PAGE gels by using Comassie blue staining, and the presence of liposomes was monitored by measuring rhodamine (Rho-PE) fluorescence in a microplate reader Synergy HT (Bio-Tek, Winooski, VT, US). Densitometry of the protein bands was performed using ImageJ software, and the percent liposome-bound protein was calculated from the band intensities measured in the third + fourth fractions (floating vesicle fractions), relative to the total sum of intensities measured in all fractions.

### Cell culture

HeLa cells stably expressing NDPK-D protein (HeLa-NDPK-D), kindly provided by Dr. Uwe Schlattner (University Grenoble Alpes, Grenoble, France), were cultured in DMEM containing 10% Fetal Bovine Serum, non-essential amino acids and 1% penicillin-streptomycin at 37°C in a humidified atmosphere with 5% CO_2_. SH-SY5Y neuroblastoma cells (ATCC^®^ CRL-2266™) obtained from Innoprot S.L. (Derio, Spain) were cultured in RIPA 1640 medium with the same supplements as described above for Hela-NDPK-D. Cells were transfected with Lipofectamine™ 3000 following the manufacturer’s instructions

### Fluorescence confocal microscopy

HeLa-NDPK-D cells were grown on 12-mm diameter glass coverslips (Paul Marienfeld GmbH & Co. KG, Lauda-Königshofen, Germany, 0111520) placed into the wells of a 24-well plate. SH-SY5Y cells were grown on μ-Slide 8 Well chambered coverslips (Ibidi, Gräfelfing, Germany, 80826). 24 h post-transfection, cells were stained with MitoTracker Red CMXRos (100 nM 1h 37 °C) and treated with 1 μM rotenone in serum-free medium (4 h for HeLa-NDPK-D and 6 h for SH-SY5Y) and then fixed with 4% PFA for 10 min. Coverslips were mounted on glass slides and samples were visualized in an inverted confocal fluorescence microscope (Nikon Eclipse C1, Nikon Inc., Melville, NY) with a x60 oil immersion objective. The excitation wavelengths used were 488 nm for GFP and 561 nm for Mitrotracker Red, and emitted fluorescence was recorded using band pass filters BP515 and BP593 respectively. Autophagy vesicles and their co-localization with mitochondria were analyzed using Just Another Colocalization Plugin Software (JaCoP) in ImageJ [**57**]. At least 30 cells were analyzed per condition.

### Statistics

Statistical analyses were performed with the Student’s t-test. Results in figure 6 were analyzed with ANOVA.

## Abbreviations

AP: Autophagosome
Atg8: Autophagy-related protein 8
CL: Cardiolipin
DMSO: Dimethyl sulfoxide
DOPE: 1,2-dioleoyl-sn-glycero-3-phosphoethanolamine
DTT: Dl-dithiothreitol
ePC: Egg phosphatidylcholine
FKBP8: Peptidyl-prolyl cis-trans isomerase FKBP8
GABARAP: Gamma-aminobutyric acid receptor-associated protein
GABARAPL1: Gamma-aminobutyric acid receptor-associated protein-like 1
GABARAPL2/GBRL2: Gamma-aminobutyric acid receptor-associated protein-like 2
GFP: Green Fluorescent Protein
IMM: Inner mitochondrial membrane
LC3A: Microtubule-associated proteins 1A/1B light chain 3A
LC3B: Microtubule-associated proteins 1A/1B light chain 3B
LC3C: Microtubule-associated proteins 1A/1B light chain 3C
LIR: LC3 interacting motif
LUV/LUVs: Large Unilamellar Vesicle/s
NDPK-D/Nm23-H4: Mitochondrial Nucleoside Diphosphate Kinase D
OMM: Outer mitochondrial membrane
PA: Phosphatidic acid
PG: Phosphatidylglycerol
PI4P: Phosphatidylinositol-4-phosphate
PINK1: PTEN induced putative kinase 1
Rho-PE: Lissamine rhodamine phosphatidylethanolamine
SDS-PAGE: Sodium dodecyl sulfate-polyacrylamide gel electrophoresis
SUV/SUVs: Small Unilamellar Vesicle/s

## Acknowledgments

The authors are indebted to Dr. C. Behrends (Munich, Germany), Dr. I. Novak (Split, Croatia), Dr. U. Schlattner (Grenoble, France), and Dr. I. Ramos (Innoprot, Derio, Spain) for providing essential plasmids and cells. They are also grateful to Ms. Araceli Marcos for skilful technical help. Ms. Maider Garnica helped in obtaining some of the mutants and performed some preliminary floating experiments. Ms. Uxue Ballesteros helped in the preparation of some of the cell experiments.

## Author contributions

AA and FMG provided the original idea, outlined the experimental approach, and obtained funding. MNI and AE designed the specific experiments, performed most of them, and analysed the results jointly with AA, LRM and FMG. YRV and JHH performed part of the experiments. MNI and AE wrote a first draft of the manuscript, with contributions from all the authors. AA reviewed the manuscript in its final form.

## SUPPLEMENTARY FIGURES

**Figure S1.**
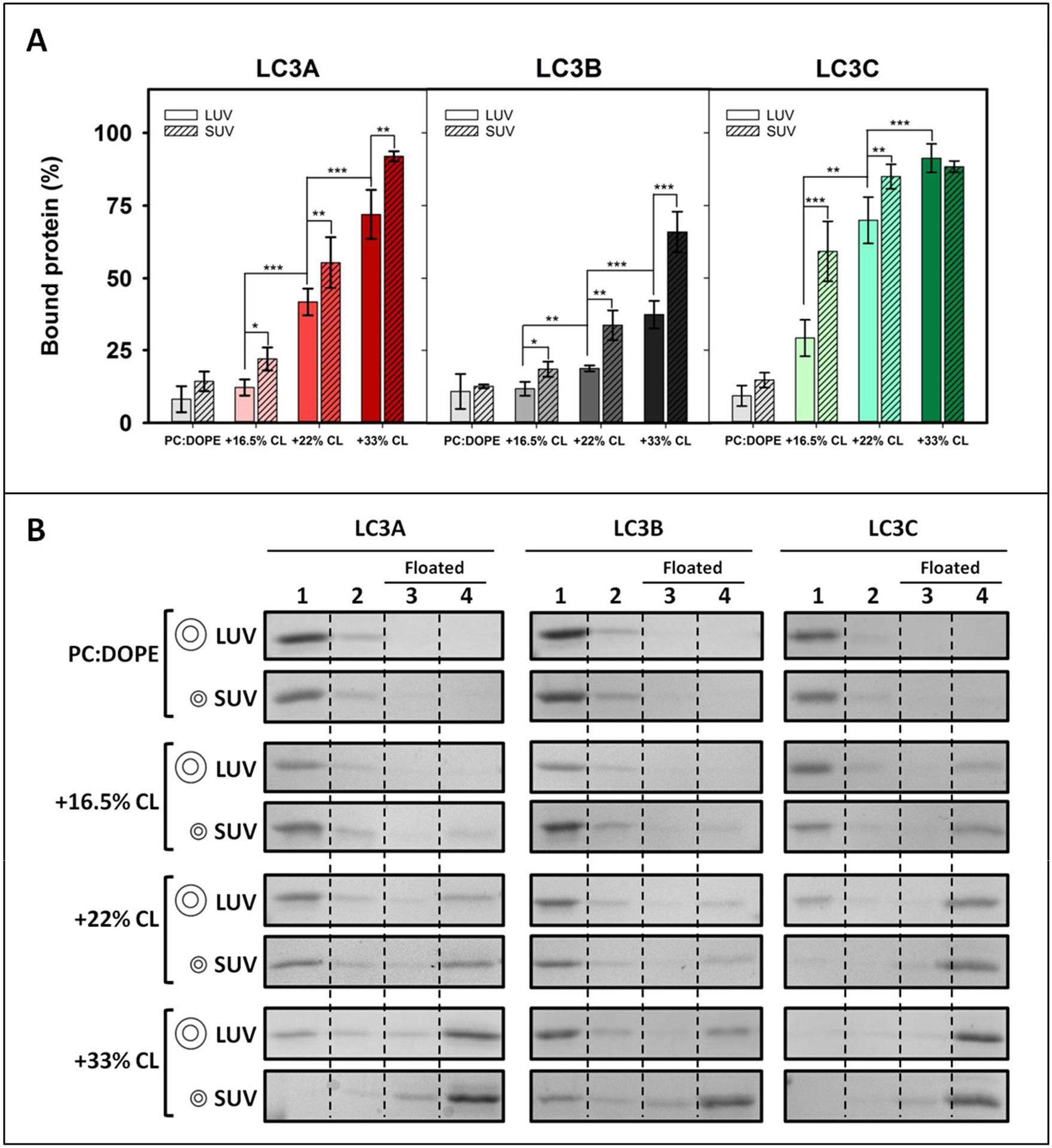
LC3 binding to liposomes is enhanced by membrane curvature and, in a dose-dependent way, by CL. Interaction of LC3 proteins with vesicles of different radii and/or CL contents was measured by a vesicle flotation assay. (**A**) Binding of LC3A, LC3B, and LC3C to liposomes, quantified by gel densitometry. LUVs (100nm) or SUVs (50nm) were composed either of ePC:DOPE (50:50 mol ratio) or of ePC:DOPE:CL. In the latter case the percent lipid proportions were 49.5:33:16.5, 44:33:22 or 33:33:33. Data shown are means ± SD (n ≥ 3). ***P < 0.001, **P < 0.01, *P < 0.05 **(B)** Representative SDS-PAGE/Coomassie Blue-stained gels of the fractions obtained from LC3A, LC3B, and LC3C vesicle flotation assays, whose quantitative results are summarized in Fig. S1A. Bound protein is computed as the proportion retrieved in fractions 3+4 (See Fig. 2A).

**Figure S2.**
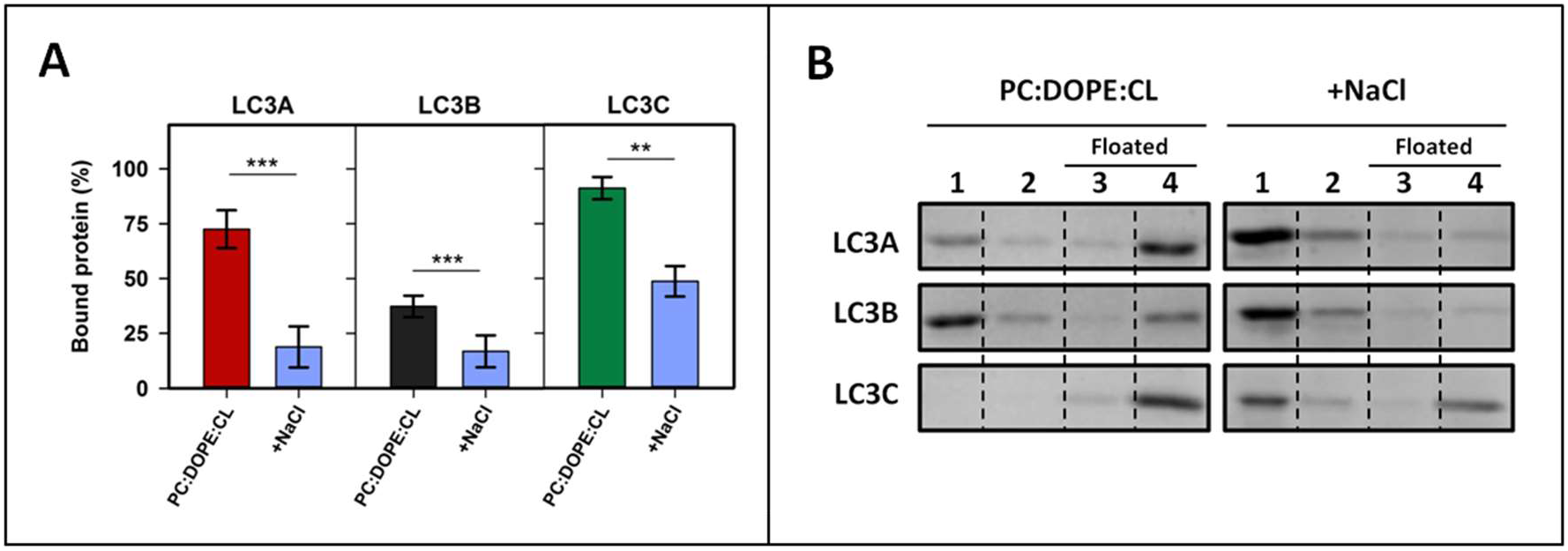
Increasing ionic strength of the medium decreases the binding of LC3 to CL. Effect of the increased ionic strength on the interaction of LC3 proteins with CL-containing membranes, measured by a vesicle flotation assay. (**A**) Binding of LC3A, LC3B, and LC3C to liposomes composed of ePC:DOPE:CL (33:33:33 mol ratio) in buffer containing 150 mM NaCl or in buffer containing 300 mM NaCl (+NaCl).Data shown are means ± SD (n ≥ 3) ***P < 0.001, **P < 0.01. (**B**) Representative SDS-PAGE/Coomassie Blue-stained gels of the fractions obtained from LC3A, LC3B, and LC3C vesicle flotation assays, whose quantitative results are summarized in Fig. S2A. Bound protein is computed as the proportion retrieved in fractions 3+4 (See Fig. 2A).

**Figure S3.**
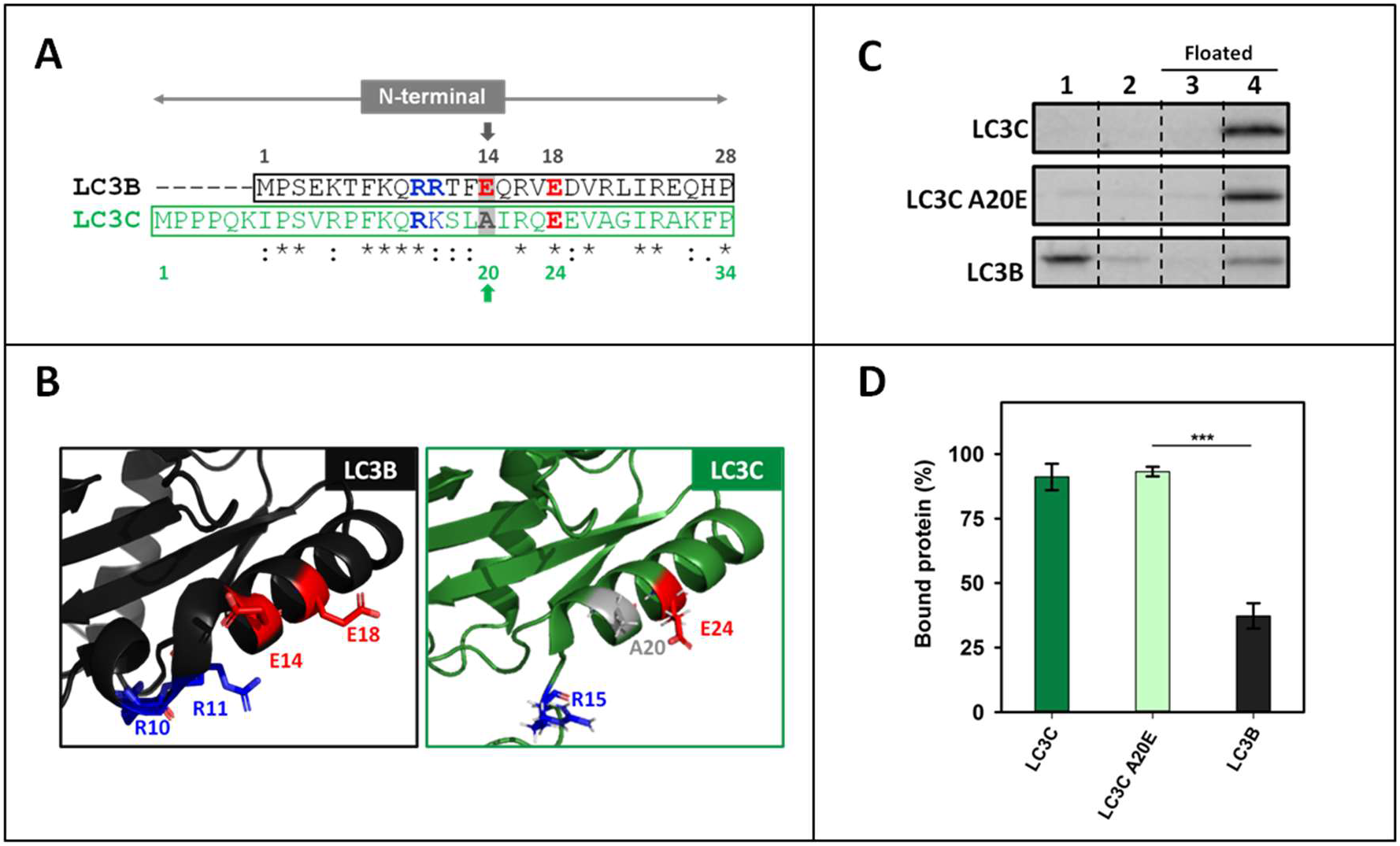
Changing a specific LC3C N-terminal residue has no effect on the protein binding to CL. (**A**) Comparative analysis of the N-terminal regions of LC3B and LC3C obtained using Clustal W. Amino acids proposed to be important in LC3B-CL interaction (R10, R11), the ones proposed to make the difference between LC3B and LC3A (E14, E18) and the corresponding ones in LC3C are colored: red, negatively-charged; blue, positively-charged; grey, no charge. The arrows show the amino acid residue designed to be mutated. (**B**) 3D structures of the N-terminal regions of LC3B and LC3C showing the said amino acid residues. (**C**) Representative SDS-PAGE/Coomassie Blue-stained gels of the fractions obtained from LC3C, LC3C A20E and LC3B vesicle flotation assays performed with CL-containing liposomes (ePC:DOPE:CL (33:33:33 mol ratio)). Bound protein is computed as the proportion retrieved in fractions 3+4 (See Fig. 2A). **(D)** Binding percentage of LC3C, LC3C A20E and LC3B to CL-containing liposomes quantified by gel densitometry. Data shown are means ± SD (n ≥ 3). ***P < 0.001.

**Figure S4.**
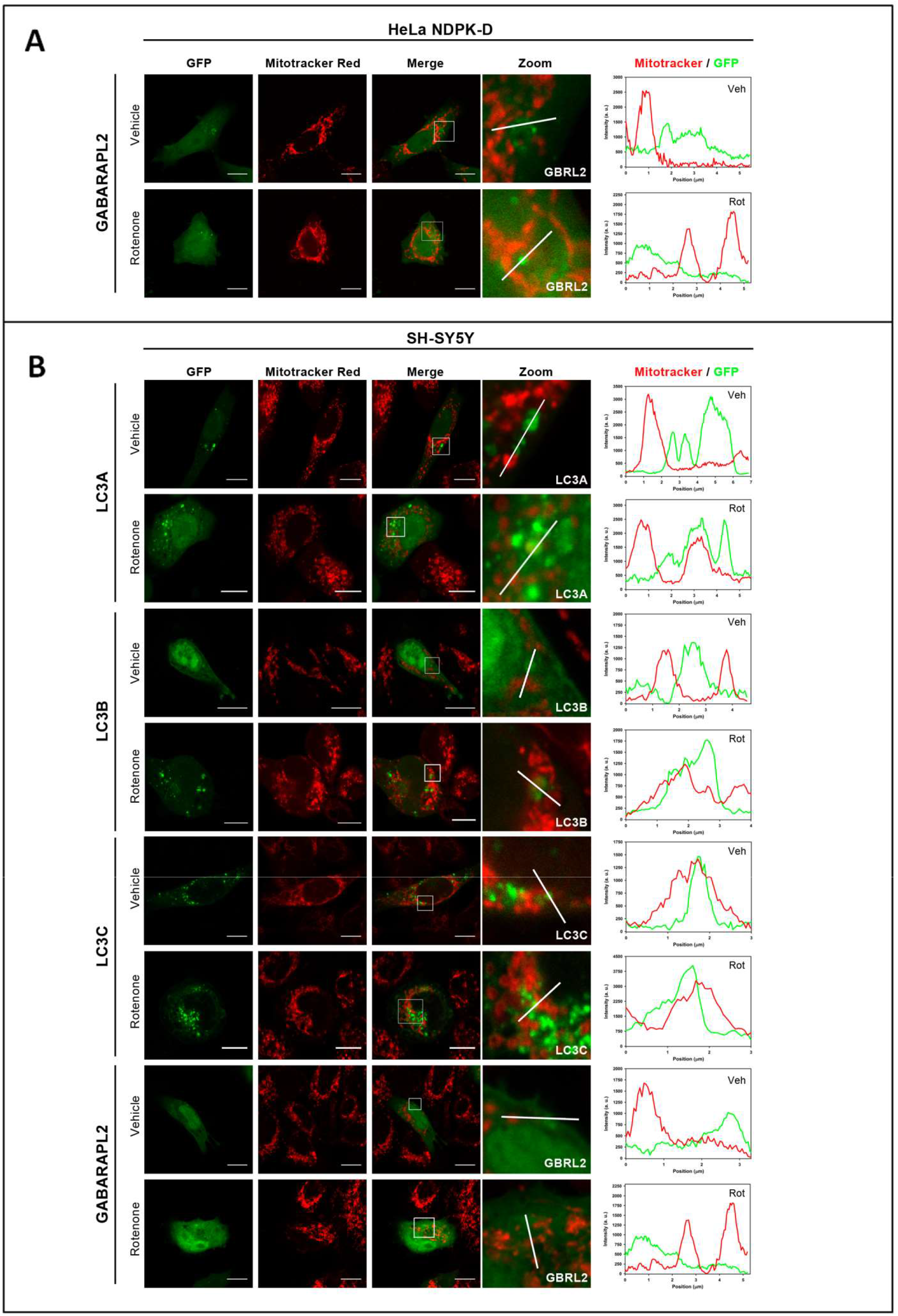
LC3A and LC3B puncta per cell and their co-localization with mitochondria increase with rotenone treatment. Cells were transfected with different members of the LC3/GABARAP family tagged with GFP. Mitochondria were labeled using Mitotracker Red, prior to treatment with 1 μM rotenone. Vehicle (Veh) controls were treated with DMSO. Images were acquired using a Nikon Eclipse C1 confocal microscope. Scale bar: 10 μm. At the right-hand side of each condition, Mitotracker (red) and GFP (green) line-profiles show examples of colocalization and non-colocalization events. (**A**) Representative images of GFP-GABARAPL2 HeLa NDPK-D transfected cells, either untreated (vehicle) or treated with rotenone. (**B**) Representative images of GFP-LC3 and GFP-GABARAPL2-transfected SH-SY5Y cells, either untreated (vehicle) or treated with rotenone.

**Figure S5.**
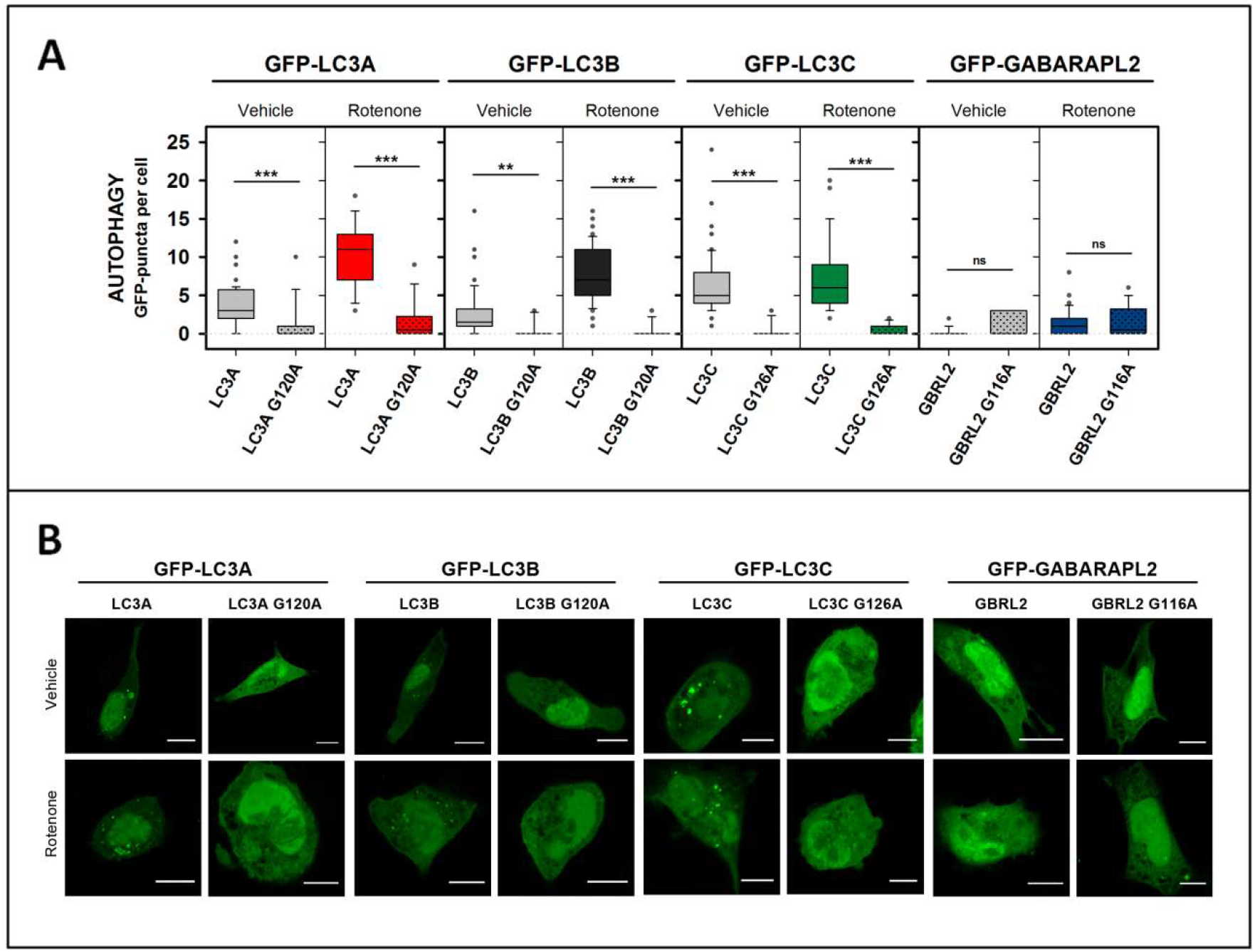
Quantification of autophagy with native LC3/GABARAP and with non-conjugatable mutants confirms that most puncta in cells are autophagic vesicles and not aggregates. SH-SY5Y neuroblastoma cells were transfected with different members of the LC3/GABARAP family tagged with GFP and their respective non-conjugatable mutant. Cells were treated with 1 μM rotenone. Vehicle (Veh) controls were treated with DMSO. (**A**) Number of GFP puncta per cell, an indication of autophagy. At least 30 images were analyzed per condition. ***P < 0.001, **P < 0.01, ns: non-significant. (**B**) Representative images of each condition. Images were acquired using a Nikon Eclipse C1 confocal microscope. Scale bar: 10 μm.

**Figure S6.**
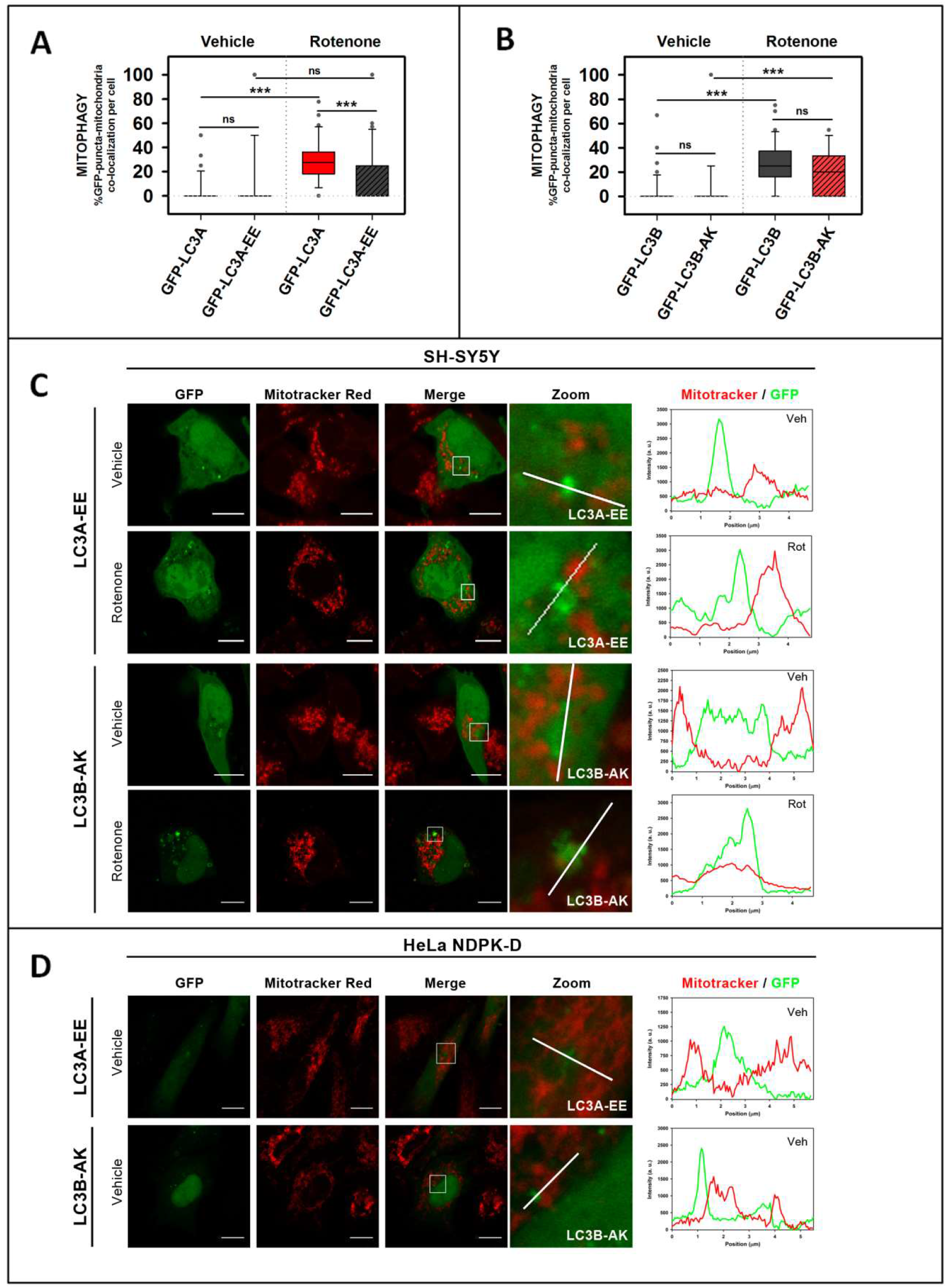
The LC3A-EE double mutation that hampers LC3A binding to CL *in vitro* also decreases its location to mitochondria in mitophagy. Cells were transfected with GFP-tagged WT or mutant LC3. Mitochondria were labeled using Mitotracker Red, prior to the treatment with 1 μM rotenone. Vehicle (Veh) controls were treated with DMSO. (**A-B**) Percent GFP puncta that colocalize with mitochondria under each condition, an indication of mitophagy. To compute the percent colocalization images were analyzed with JACop plugging of Image J. At least 30 images were analyzed per condition. ***P < 0.001, ns: non-significant. (**C**) Representative images of GFP-LC3A-EE- and GFP-LC3B-AK-transfected SH-SY5Y cells, either untreated (vehicle) or treated with rotenone. (**D**) Representative images of GFP-LC3A-EE- and GFP-LC3B-AK-transfected, rotenone-untreated HeLa NDPK-D cells. Images were acquired using a Nikon Eclipse C1 confocal microscope. Scale bar: 10 μm. At the right-hand side of each condition, Mitotracker (red) and GFP (green) line profiles show examples of colocalization and non-colocalization events.

**STable 1:**
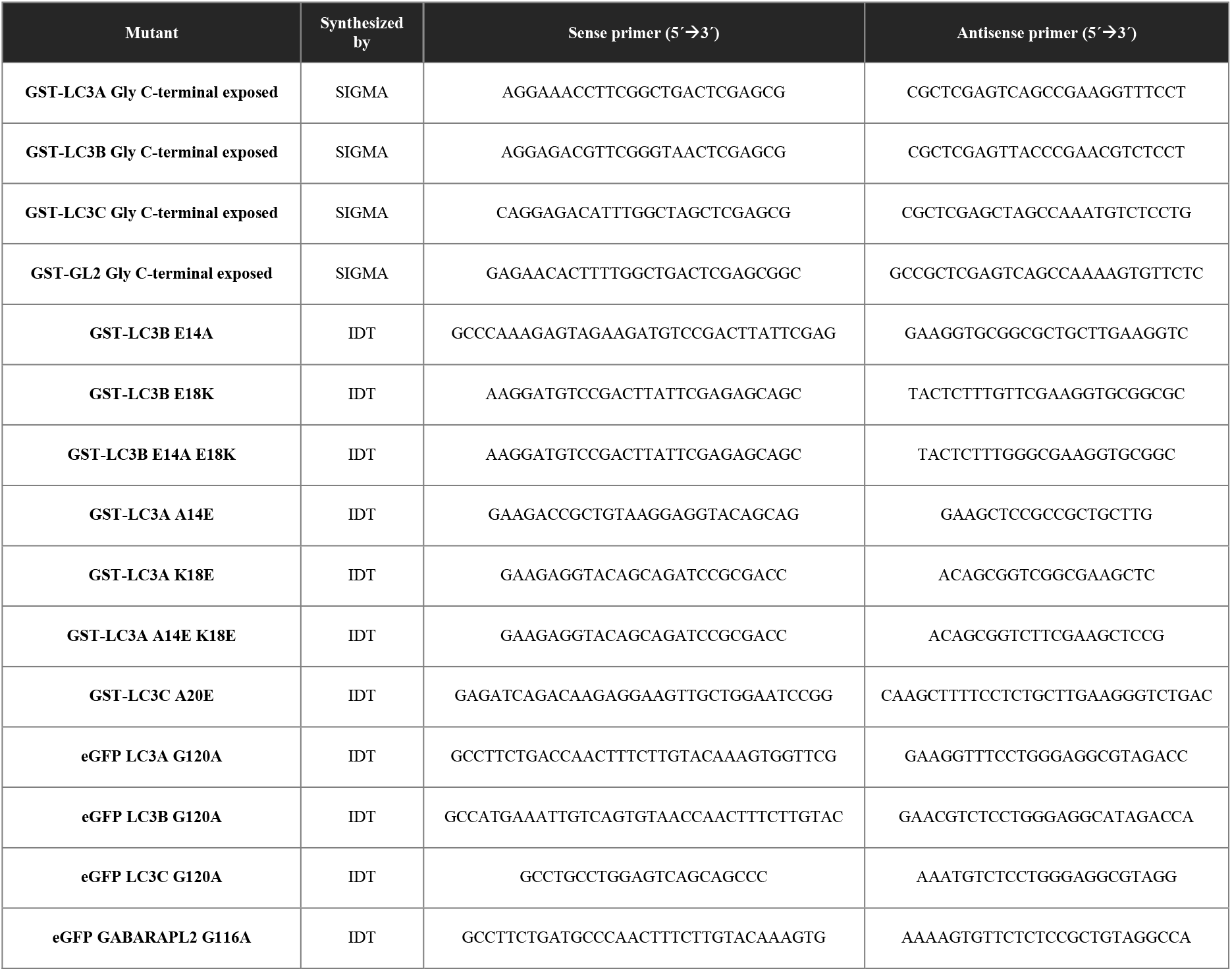
Primers used to perform site-directed mutagenesis

